# Identification of Immune-Related Candidate Biomarkers in Plasma of Patients with Sporadic Vestibular Schwannoma

**DOI:** 10.1101/2023.01.24.525436

**Authors:** Sasa Vasilijic, Nadia A. Atai, Hiroshi Hyakusoku, Steven Worthington, Yin Ren, Jessica E. Sagers, Mehmet I Sahin, Takeshi Fujita, Lukas D. Landegger, Richard Lewis, D. Bradley Welling, Konstantina M. Stankovic

## Abstract

Vestibular schwannoma (VS) is intracranial tumor arising from neoplastic Schwann cells, causing hearing loss in about 95% of patients. The traditional belief that hearing deficit is caused by physical expansion of the VS, compressing the auditory nerve, does not explain the common clinical finding that patients with small tumors can have profound hearing loss, suggesting that tumor-secreted factors could influence hearing ability in VS patients. Here, we conducted profiling of patients’ plasma for 67 immune-related factors on a large cohort of VS patients (N>120) and identified candidate biomarkers associated with tumor growth (IL-16 and S100B) and hearing (MDC). We identified the 7-biomarker panel composed of MCP-3, BLC, S100B, FGF-2, MMP-14, eotaxin, and TWEAK that showed outstanding discriminatory ability for VS. These findings revealed possible therapeutic targets for VS-induced hearing loss and provided a unique diagnostic tool that may predict hearing change and tumor growth in VS patients and may help inform the ideal timing of tumor resection to preserve hearing.

**Teaser:** Profiling of plasma in vestibular schwannoma patients revealed biomarkers that could predict hearing change and tumor growth.

## Introduction

Vestibular schwannoma (VS) is an intracranial tumor arising from neoplastic Schwann cells in the vestibular branch of the eighth cranial nerve (*1*). VS is primarily caused by mutations in the neurofibromin 2 (*NF2*) gene disrupting production of merlin, a tumor suppressor. *NF2* mutations may be tumor-specific, as in unilateral, sporadic VS, or rare germline mutations leading to bilateral VS and *NF2*-related schwannomatosis (US incidence 1:10,000 vs. 1:33,000, respectively) (*2–4*). The most common symptoms of VS are sensorineural hearing loss and tinnitus (95% and 60% of patients, respectively), followed by balance dysfunction (*5, 6*). Although histologically non-malignant, VS can cause substantial morbidity due to where it typically arises within the internal auditory canal, and can be life-threatening if unchecked expansion into the cerebellopontine angle compresses the brainstem (*7*).

VS management is informed by the rate of tumor growth, which is monitored with clinical imaging (i.e., magnetic resonance imaging [MRI]) (*8*). Growing tumors are typically treated with microsurgical resection or stereotactic radiation therapy. However, the size and location of sporadic VS do not necessarily correlate with the severity of the tumor-associated hearing loss (*9*), thus, the “wait and scan strategy” may be insufficient to determine the ideal timing for tumor resection to prevent progressive hearing loss (*10–12*).

The identification of blood biomarkers whose levels correlate with hearing loss severity or tumor size, or are prognostic of clinical outcomes, could have immense value for the management of VS (*13*). Prior attempts to characterize potential blood biomarkers have focused on NF2-related rather than the more common sporadic VS, which comprises >90% of cases (*2, 3*) and had limited sample sizes (<30 patients) (*14–16*). Therefore, we conducted extensive immune profiling of blood plasma to identify candidate biomarkers of the tumor and related hearing loss in sporadic VS, informed by our previous studies *in vitro* or among smaller cohorts. For example, matrix metalloproteinase 14 (MMP-14) is the most abundant matrix metalloproteinase in VS, with differential expression between tumors associated with poor hearing (PH) versus good hearing (GH) (*17*). Additionally, sporadic VSs associated with GH secreted high levels of fibroblast growth factor 2 (FGF-2), which had an otoprotective effect *in vitro (18*). Conversely, higher secreted levels of tumor necrosis factor alpha (TNF-α) were associated with worse VS-induced hearing loss and had an ototoxic effect *in vitro (19*). Finally, interleukin 18 (IL-18) levels were significantly elevated in the tumors of VS-PH patients compared to VS-GH patients (*20*).

Motivated by these data, we quantified the levels of MMP-14, FGF-2, IL-18, and TNF-α in the plasma >120 patients with sporadic VS and extended the analysis to 67 cytokines, chemokines, growth factors, and cell surface proteins (**Table S1**). We identified and analyzed candidate biomarkers with differential concentrations in the plasma of patients with and without VS (controls) and examined their association with pre-operative hearing and tumor volume. Finally, we assessed the diagnostic utility of the candidate biomarkers and a composite biomarker panel in discriminating between VS patients and controls.

## Results

### Patient characteristics

A total of 163 patients with sporadic VS, including 34 with GH and 124 with PH (Fig. S1), were included for comparison with 70 controls. VS patients and controls had similar mean ages (56 vs. 46 years) and proportion of females (55-56%); VS-PH patients were significantly older than controls (53 years; *P*=0.002) (Fig. 1). Compared with VS-GH patients, VS-PH patients had significantly larger tumor volume (7.39 vs. 3.38 cm^3^), worse ipsilateral pure tone average (PTA; 62.37 vs. 17.33 dB) and word recognition percentage (WR; 39.20% vs. 94.21%), and worse contralateral PTA (19.75 vs. 8.58 dB; all *P*<0.005) (Fig. S2).

**Fig 1.**
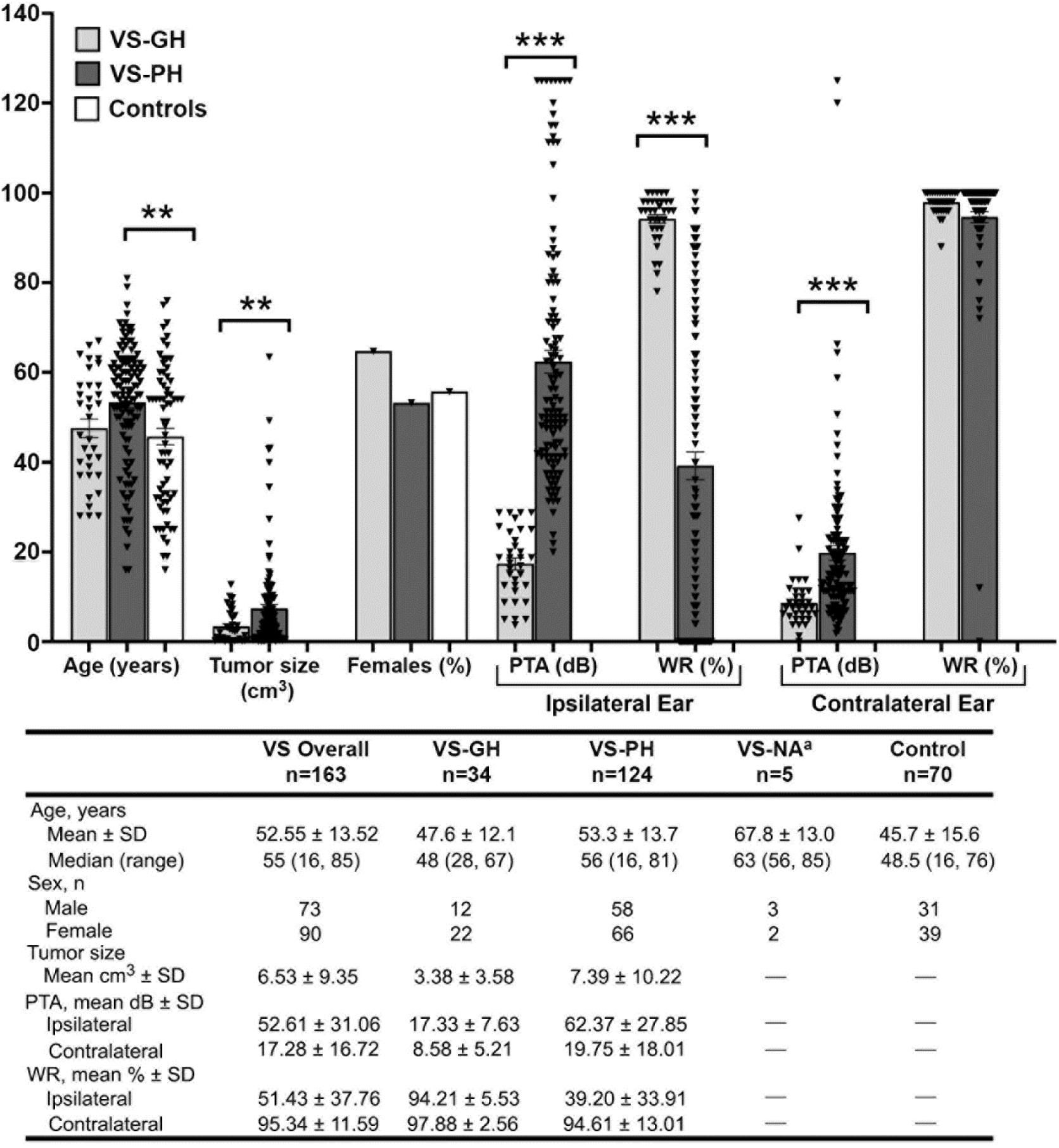
Demographic and clinical characteristics of the VS and control cohorts. Abbreviations: dB, decibel; NA, not available; SD, standard deviation; VS, vestibular schwannoma; VS-GH, vestibular schwannoma patients with good hearing; VS-PH, vestibular schwannoma patients with poor hearing. ** p<0.01, *** p<0.001. Note: a NA refers to the small group of patients with poor hearing (n=5) who did not have quantitative audiograms available.

### Plasma levels of candidate biomarkers

#### VS patients vs. controls

Twenty-two of the 67 profiled factors were detectable in the plasma of >75% of VS patients and considered candidate biomarkers (**Table S2**). In both VS and control patients, the highest mean plasma values were for MMP-14, IL-2R, and SDF-1α, while the lowest values were for MCP-2, eotaxin, and MCP-3. The levels of candidate biomarkers measured in controls are in agreement with prior reports (*21–25*). The levels of 16 candidate biomarkers significantly differed between the VS and control groups (Fig. 2a). The most elevated factors among VS patients vs. controls were MMP-14 and FGF-2 (both *Padj<0.001*), where the ratio of plasma levels were ~7.5 and 4 times higher, respectively (Fig. 2b). MMP-14 was ~2-fold higher, while IL-18 was ~2-fold lower, in VS patients vs. controls (Fig. 2c). TNF-α was detected at very low levels in only 24% of analyzed patients and was omitted from the analysis (**Table S2**).

**Fig 2.**
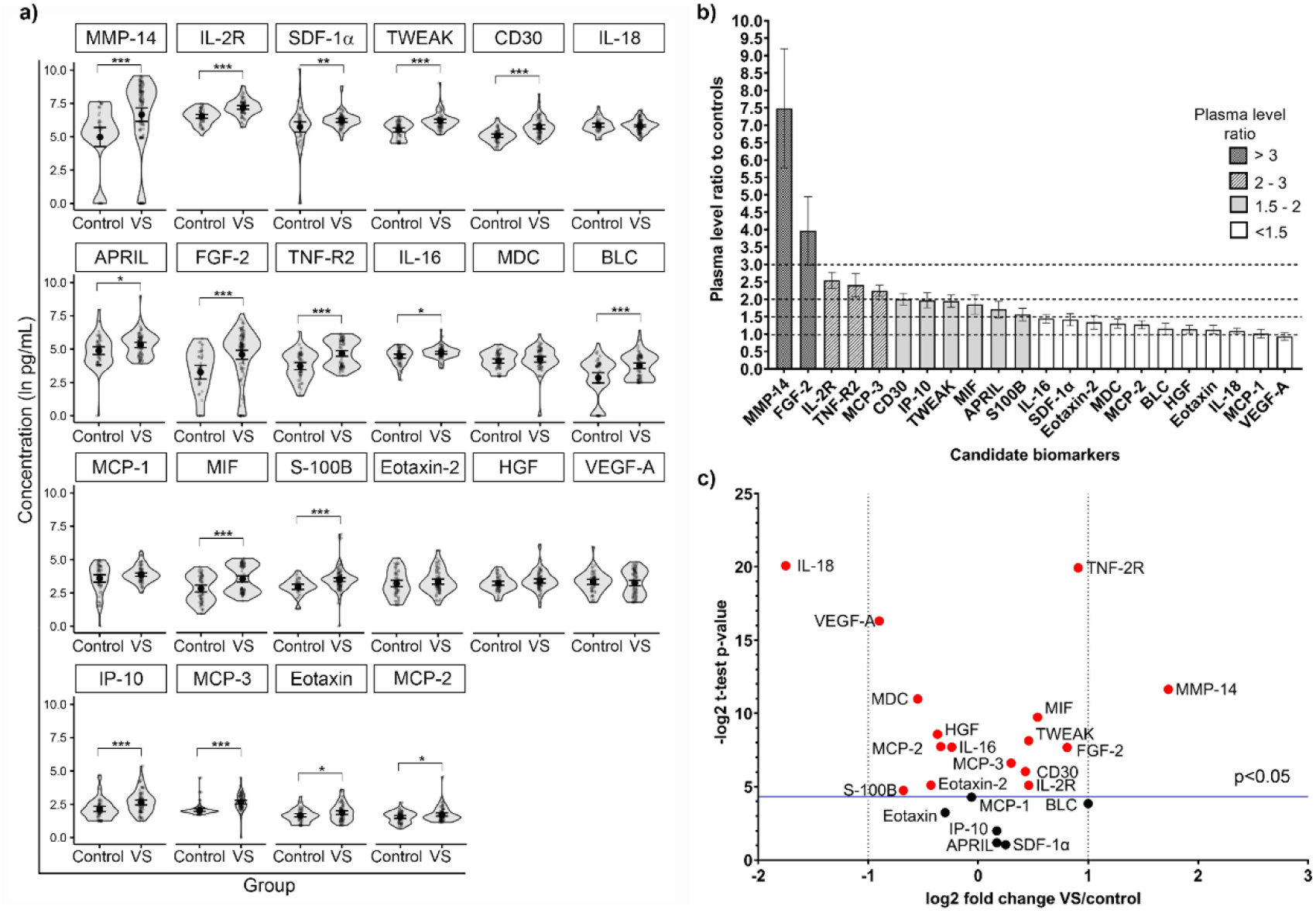
Comparison of candidate biomarker plasma levels between VS patients and controls. (a) The concentrations of 16 candidate biomarkers significantly differed between VS patients and controls. *Padj<0.05, **Padj<0.01, ***Padj<0.001. Ratios (b) and fold change (c) of candidate biomarker levels between VS patients and controls. In (c), the blue line indicates the threshold for significance. The full names of candidate biomarkers are listed in S1 Table. Abbreviations: VS, vestibular schwannoma.

Ten biomarkers (MCP-3, CD30, IL-2R, TWEAK, TNF-R2, S100B, FGF-2, MIF, BLC, and IP-10) were significantly elevated among VS patients of both sexes compared to controls (all *Padj<0.05*) (Fig. S3). Eotaxin, MCP-2, MMP-14, and SDF-1α were significantly elevated only in male VS patients while APRIL was only elevated in female VS patients (all *Padj<0.05*).

#### VS-GH and VS-PH patients vs. controls

Compared to controls, 11 candidate biomarkers (MCP-3, CD30, S100B, TNF-R2, TWEAK, IL-2R, MIF, BLC, IL-16, SDF-1α, and IP-10) were significantly elevated in both VS-GH and VS-PH patients, while FGF-2, MMP-14, APRIL, and MCP-1 significantly differed only in VS-PH patients (Fig. S4). Controlling for sex and age, VS-PH patients had the highest increases of FGF-2 (43.96%) and MMP-14 (37.91%), while VS-GH patients had the highest increases of MCP-3 (45.10%) and BLC (39.50%).

Compared to controls, there were significant sex simple effects among VS-PH patients of both sexes for levels of CD30, FGF-2, IL-2R, IP-10, MCP-3, MIF, S100B, TNF-R2, and TWEAK; and among males only for BLC, eotaxin, MCP-1, MCP-2, MMP-14, and SDF-1α (*Padj<0.05*) (Fig. S5a). There were significant sex simple effects among VS-GH patients of both sexes vs. controls for IL-2R and MCP-3; among females only for CD30, S100B, and TWEAK; and among males only for MIF, SDF-1α, and TNF-R2 (all *Padj<0.05*) (Fig. S5b).

#### VS-GH vs. VS-PH patients

The level of IL-16 was significantly higher in VS-GH vs. VS-PH patients, and MCP-3 had the highest significant percent change between groups when controlling for sex, age, and tumor volume (both *Padj<0.05*) (Fig. S4).

### Interaction of biomarker levels with pre-operative hearing

The relationships between candidate biomarker levels and pre-operative PTA and WR were assessed among VS patients with a robust linear model (RLM) and a fractional logit model (FLM), respectively. There were no significant associations with PTA and biomarker levels, or sex-specific effects (Fig. S6, S7a). However, there was a significant association with ipsilateral WR scores and MDC levels (*Padj=0.021*), while MCP-3 approached significance (*Padj=0.054*) (Fig. 3). MCP-3 had the greatest effect according to the odd ratio calculated for the candidate biomarkers positively associated with WR, where a 1-unit increase in its natural log was associated with a WR increase of 135.52% (vs. +89.09% for MDC). Additionally, significant simple sex effects with WR were observed among males for IL-16, MCP-3, and MDC (all *Padj* <0.05) (Fig. S7b), and the odds ratio indicated that a 1-unit increase in the natural log of IL-16 would have a greater impact on WR (+1108.69%) than MCP-3 (+360.79%) or MDC (+129.27%).

**Fig 3.**
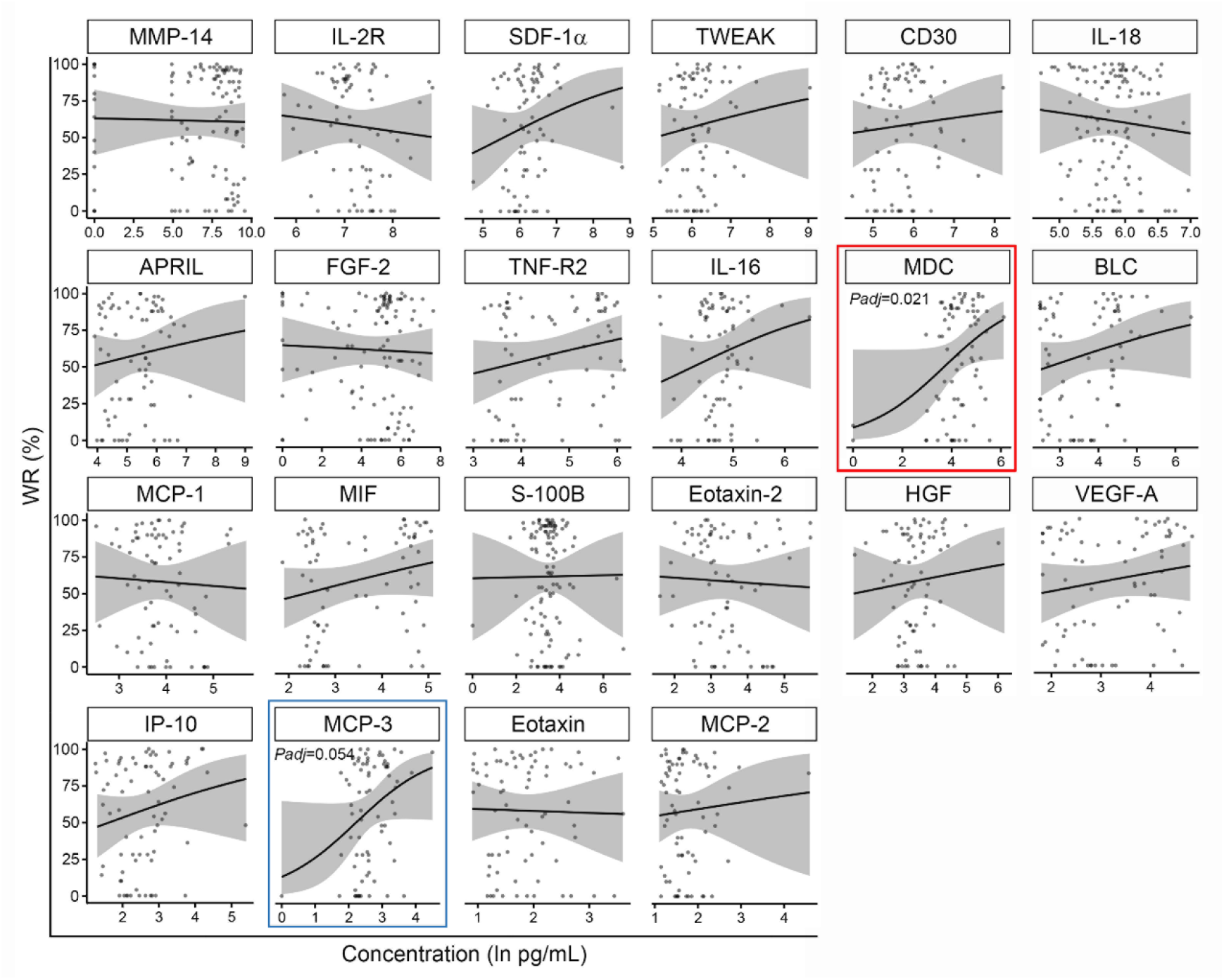
Plasma level of candidate biomarkers and pre-operative word recognition score in VS patients. The red box indicates a significant association between MDC levels and pre-operative WR; the blue box indicates an association approaching significance for MCP-3. Abbreviations: VS, vestibular schwannoma; WR, word recognition.

### Interaction of biomarker levels with pre-operative tumor volume

In the RLM assessing the relationship of biomarker levels and tumor volume, significant associations were found for IL-16 and S100B (Fig. 4), and there was a significant simple sex effect among females for S100B and tumor volume (all *Padj<0.05*) (Fig. S8).

**Fig 4.**
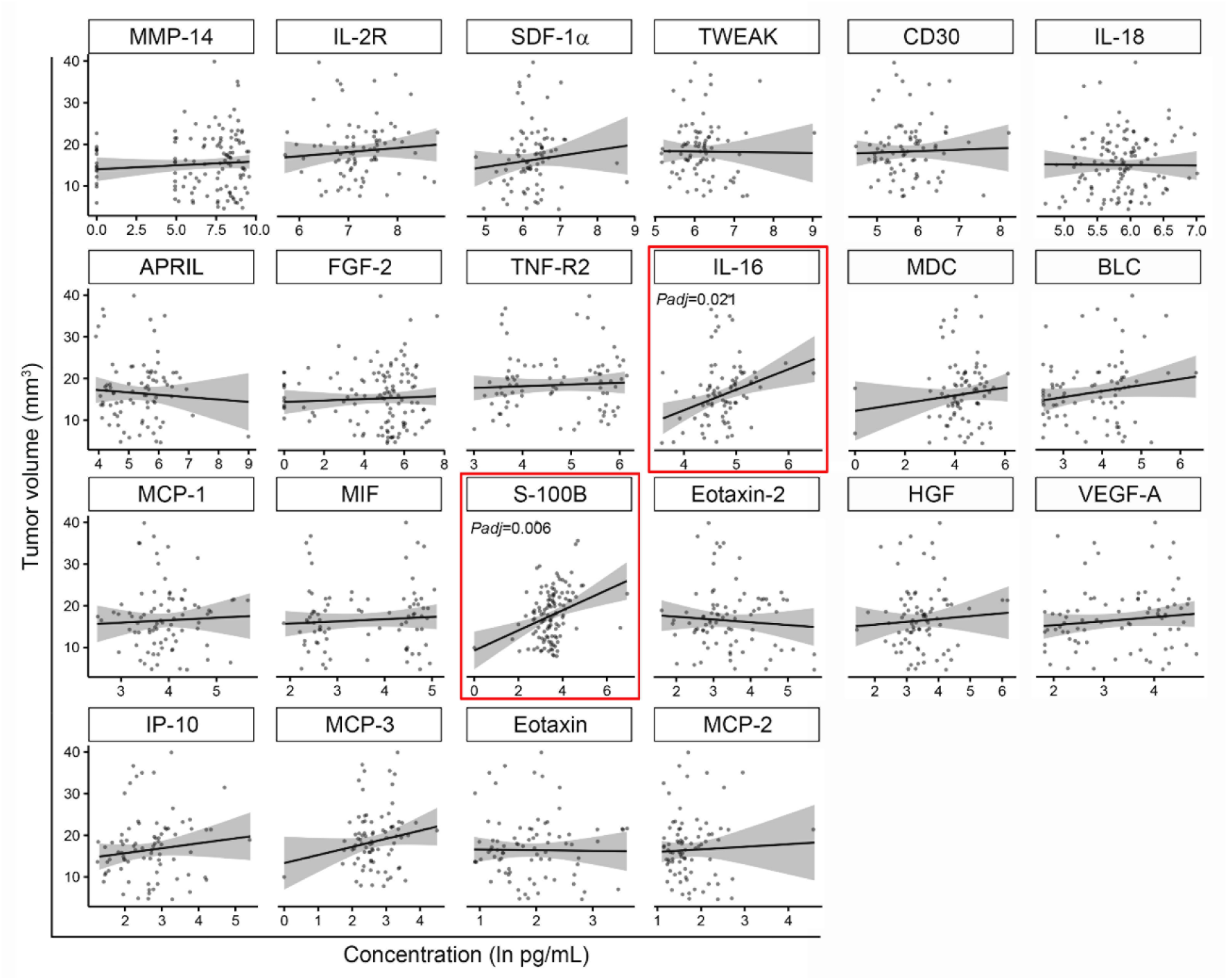
Plasma levels of candidate biomarkers and tumor volume in VS patients. The red boxes indicate significant associations between IL-16 and S100B levels and pre-operative tumor size. Abbreviations: VS, vestibular schwannoma.

### Discriminative power, diagnostic utility, and relationships of candidate biomarkers

In the receiver-operating characteristic curve (ROC) analysis of the balanced dataset of sex- and age-matched VS patients and controls, 12 of the 16 significantly elevated factors in VS patients’ plasma had area under the curve (AUC) values >0.7 and were considered potentially predictive (Fig. 5). AUC values were outstanding (0.9-1) for TNF-R2 and MIF; excellent (0.8-0.9) for CD30, MCP-3, IL-2R, BLC, and IP-10; acceptable (0.7-0.8) for MMP-14, TWEAK, eotaxin, FGF-2, and S100B; and poor (0.5-0.7) for APRIL, IL-16, SDF-1α, and MCP-2.

**Fig 5.**
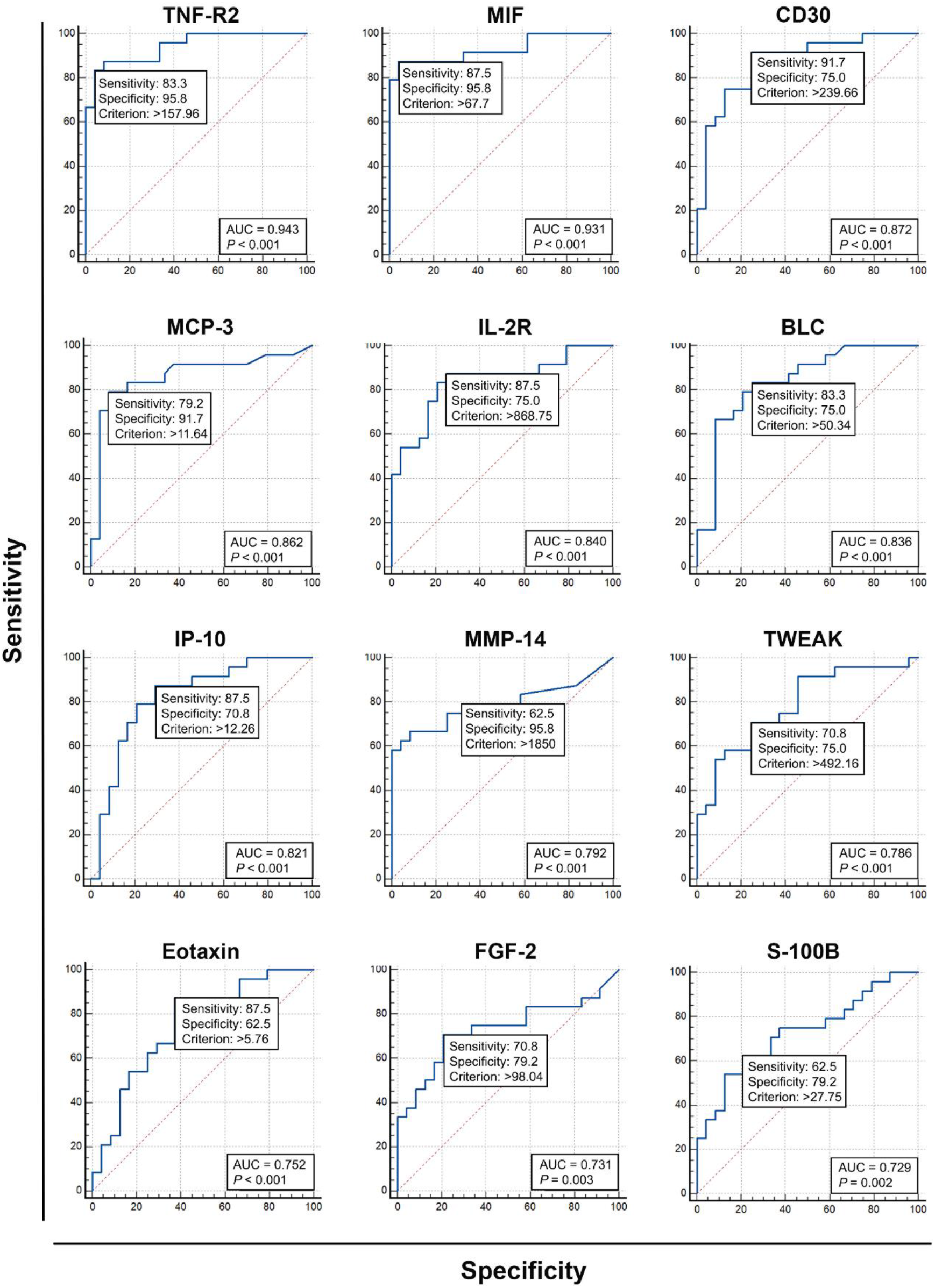
Diagnostic utility and discriminatory power of candidate biomarkers with significant AUC values. Abbreviations: AUC, area under the curve.

To evaluate whether a biomarker panel could discriminate between VS patients and controls, 7 biomarkers (MCP-3, BLC, S100B, FGF-2, MMP-14, eotaxin, and TWEAK) were selected based on their predictive power and correlation with hearing or tumor size. A combinatorial analysis of the 7 biomarkers indicated 120 different panels, of which 39 (32.5%) had an outstanding discrimination level (**Table S3**). The 7-biomarker panel demonstrated the best predictability, reaching an AUC of 0.934 with 87.5% sensitivity and 95.8% specificity. This was a 19.13% improvement compared to the mean AUC of the individual biomarkers (AUC7-panel: 0.934 vs. AUCmean of 7: 0.784). This was confirmed by logistic regression analysis (**Table S4)**, and the accuracy and utility of the 7-biomarker panel were verified by performing 10-fold cross-validation and permutation tests (Fig. 6).

**Fig 6.**
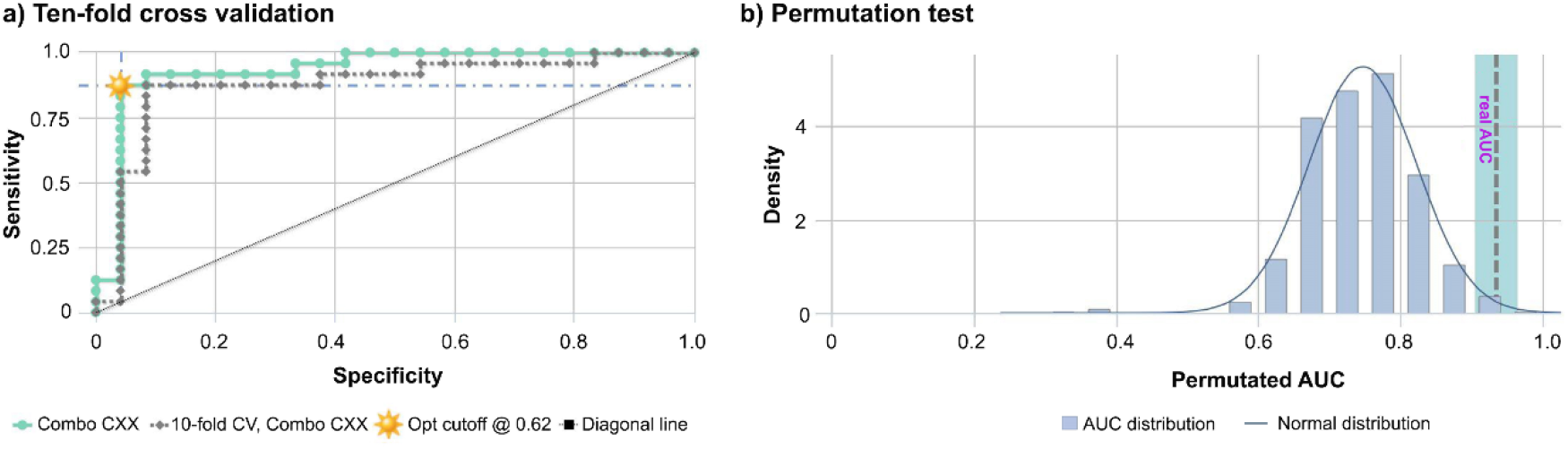
Ten-fold cross-validation and permutation tests of the panel of 7 candidate biomarkers selected as the best predictors associated with hearing and tumor size in VS patients. The 7 biomarkers in the panel (CCX) were MCP-3, BLC, S100B, MMP-14, TWEAK, FGF-2, and eotaxin. Abbreviations: AUC, area under the curve; CCX, biomarker combination 120 of 120; CV, cross-validation; Opt, optimal; VS, vestibular schwannoma.

Of the 39 panels with outstanding predictability, MCP-3 and eotaxin were the most commonly appearing biomarkers (100% and 79.5% of panels, respectively). BLC, S100B, FGF-2, MMP-14, and TWEAK participated in 46-61% of panels. Analyzing the area under the precision recall curve (AUPRC) revealed better classifier performance for MCP-3 vs. BLC (AUPRC: 0.859 vs. 0.812), MMP-14 vs. FGF-2 (0.860 vs. 0.791), and TWEAK vs. eotaxin (0.808 vs. 0.724) (Fig. S9).

A mutual comparison of the 7 candidate biomarkers with Spearman correlation analysis resulted in 23 correlation pairs. VS patients had more significant correlations than controls (12 vs. 4), suggesting a disease-specific relationship between candidate biomarker levels in ~50% of correlation pairs. Both VS patients and controls had significant positive correlations between 1) MCP-3 and BLC; 2) MCP-3 and MMP-14; and 3) BLC and MMP-14. However, correlations between 1) BLC and FGF-2/S100B/TWEAK; 2) MMP-14 and FGF-2/S100B/eotaxin; 3) MCP-3 and FGF-2; and 4) S100B and FGF-2/TWEAK were significant only in VS patients (all *P*<0.05) (Fig. S10). The strongest positive correlation in VS patients was between MMP-14 and FGF-2; there were moderate positive correlations between MCP-3 and BLC, TWEAK and BLC, and FGF-2 and S100B (all *P*<0.05). Eotaxin, MCP-1, and IP-10 levels were negatively correlated with MMP-14 levels in VS patients (all *P*<0.05). In controls, significant correlations were identified between MCP-3 and MMP-14, MCP-3 and TWEAK, MCP-3 and BLC, BLC and MMP-14 (all *P*<0.05).

## Discussion

In this largest immune profiling of sporadic VS patients to date, we identified several groups of potential circulating biomarkers. IL-16 and S100B were significantly elevated across all VS groups (overall, VS-GH, and VS-PH) vs. controls and associated with WR and/or larger tumor volume. MDC was not elevated in any VS group vs. controls but was associated with better WR score. Furthermore, MCP-3, BLC, SDF-1α, MMP-14, and MIF were elevated in specific VS group(s) vs. controls and showed a trend of association with GH and/or tumor volume. The remaining factors were elevated in specific VS groups but not associated with tumor size or hearing: CD30, IL-2R, IP-10, and TNF-R2 were significantly elevated across all VS groups; FGF-2, APRIL, MCP-2, and eotaxin were significantly elevated in VS and VS-PH patients; and MCP-1 was only elevated in VS-PH patients.

A possible explanation for the positive relationship between IL-16 and good pre-operative WR could be the neuroprotective effects reported for neuronal IL-16 in reducing kainate-induced excitotoxicity and facilitating neurite outgrowth (*26–29*). IL-16 may also be considered a tumor growth-related biomarker given its association with larger tumor volume in VS patients, consistent with prior studies noting a correlation between IL-16 and tumor progression in other disorders (*30*). As the pro-IL-16 form acts as a cell cycle suppressor, mature IL-16 generated by proteolytic activity of caspase-3 is more likely to be involved in VS tumor growth (*31*). Mature IL-16 can stimulate chemotaxis and proliferation of T and B cells and the recruitment of pro-tumorigenic macrophages (*31*).

The significant elevation in the levels of the proinflammatory chemokine MCP-1 in VS-PH patients vs. controls, particularly among males, suggests a relationship with poor pre-operative hearing. A similar effect has been found in Meniere’s Disease, where higher plasma MCP-1 values were associated with progression of hearing impairment and bilateral disability (*32*). Although the pathophysiological mechanism of MCP-1-mediated hearing loss is unknown, MCP-1 could amplify cochlear inflammation by recruitment of monocytes and macrophages (*33*), which may then produce proinflammatory, ototoxic factors such as TNF-α (*19, 34*). The upregulation of MCP-1 may be due to the lack of merlin in VS, which normally suppresses NF-κB signaling (*35–37*). A significant sex-interaction effect was not observed for MCP-1, although it was significantly elevated only among male VS-PH patients and showed a trend of positive association with higher PTA values. Estradiol has been reported to inhibit ongoing autoimmune neuroinflammation and NFκB-dependent MCP-1 expression in reactive astrocytes (*38*),thus a similar process may be at play in VS where lack of estradiol contributes to higher MCP-1 levels and potentially poorer hearing in males.

Schwann and immune cells, including macrophages, express hormone receptors which can promote immune response or VS tumor growth in sex-specific directions (*39–45*). For example, macrophages from males produce higher levels of proinflammatory cytokines while those from females produce more anti-inflammatory IL-10 and have higher phagocytotic activity (*46*). The significant association of larger tumor size among female VS patients with S100B, a Schwann cell and astrocyte marker, is notable given ongoing interest in S100B as a prognostic factor in other disorders (*47–50*). Serum S100B concentration and VS tumor size are correlated among both males and females with presumed sporadic VS (*51*), but not with NF1, NF2, or schwannomatosis (*14*). Thus, the prognostic value of S100B may depend on VS tumor etiology as well as patient sex. Our results also suggest a sex-dependent relationship between higher levels of macrophage regulator chemokines (IL-16, MCP-3, and MDC) and good pre-surgical WR among male VS patients. IL-16 has a role in macrophage polarization, with higher expression promoting the M1 phenotype (*52*), while MCP-3 and MDC attract and activate multiple immune cells, including monocytes (*53*).

Finally, we demonstrated that a 7-biomarker panel had outstanding discrimination ability for VS. MMP-14 and FGF-2 were chosen as two highly elevated biomarkers in VS patients with strong disease-specific positive intercorrelations, MCP-3 and BLC had positive correlations with GH, S100B is a phenotypic marker of VS correlating with tumor size, and TWEAK and eotaxin were selected based on their contribution to improving the predictability (panel AUC) of the other biomarkers. Correlation analysis showed significant interconnections between the panel biomarkers, suggesting possible roles in VS pathogenesis. Overall, our findings suggest that the 7-biomarker panel could be an additional tool to monitor or predict hearing change and tumor growth in VS patients, and help inform diagnoses or the ideal timing of tumor resection to preserve hearing.

One limitation of our study is that it is cross-sectional and therefore the temporal link between the outcome and VS presence cannot be determined because both are examined simultaneously. A future longitudinal study focused on the validation of our findings is required. Such a study may benefit from multi-institutional patient accrual because VS is a rare tumor.

In summary, this is the first study conducting robust immune profiling of blood plasma from a large cohort of patients with sporadic VS for comparison with controls, between VS-GH and VS-PH patients, and between sexes. We correlated biomarker levels with PTA, WR, and tumor size and validated a potentially diagnostic 7-biomarker composite panel with outstanding discriminatory ability for VS.

## Materials and Methods

### Study population and specimen collection

From 07/2015–04/2021, blood was prospectively collected from patients undergoing VS resection at Massachusetts Eye and Ear Infirmary (MEEI) in Boston, MA on the day of surgery, typically within 30 minutes of inducing general anesthesia and ≥1 hour before tumor microdissection. Blood from controls was collected at the Massachusetts General Hospital (MGH) Blood Donor Center in Boston, MA, and the Research Blood Components in Watertown, MA. At collection, fresh blood was stored in EDTA vacutainer tubes (Becton Dickinson, NY, US) and kept at 4°C without freezing. The whole blood samples were centrifuged at 2000 g for 10 min at 4°C. Plasma was separated and spun at 2000 g for 5 min at 4°C. Centrifuged plasma was filtered through 0.8 μm filter units (MF-Millipore MCE membrane, SLAA033SB; Millipore, Burlington, MA, US) and stored at −80°C until further use.

Eligible patients had unilateral, sporadic VS that had not been previously resected or irradiated. Of 186 enrolled VS patients, 163 met these criteria and were included in the analyses (CONSORT diagram in **Fig S1**) for comparison with 70 controls.

### Clinical data

Clinical and demographic data were collected from patient charts, operative reports, pathology reports, and pre-operative radiographic imaging. Patient variables included age at tissue collection; pre-surgical tumor volume measured via high-resolution axial contrast-enhanced T1-weighted brain MRI; internal auditory canal protocol (*54*); and pre-surgical pure-tone audiometric threshold and WR measurements. MRI and hearing tests were those nearest to resection, typically ≤3 months.

WR was defined as the percentage of spoken monosyllabic words discernable from a list typically read at 70 dB or the level at which a patient’s speech intelligibility curve plateaus. Pure-tone audiometric thresholds at 0.5, 1, 2, and 3 kHz were used to calculate the PTA. GH was defined as WR>70% and PTA<30 dB per the American Academy of Otolaryngology–Head and Neck Surgery criteria (*55*). Otherwise, patients were classified as having PH. A deaf ear was assigned a PTA of 125 dB and WR score of 0%.

### Biomarker measurements

Luminex (65 cytokines, chemokines, growth factors, and soluble receptors); electrochemoluminescence (IL-18, TNF-α, and FGF-2); and ELISA (S100B and MMP-14) assays were conducted as described in detail below. Potential biomarkers were defined as those detectable in the plasma of >75% of patients. The list of analyzed biomarkers is in **Table S1.**

#### Luminex assay

Simultaneous multiplex profiling of 65 immuno-related factors composed of cytokines, chemokines, and growth factors was performed using a customized multiplex bead-based immunoassay - Immune Monitoring 65-Plex Human ProcartaPlex™ Panel (#EPX650-10065-901; ThermoFisher Scientific, Waltham, MA, US) according to manufacturer’s instructions. The fluorescence-based signal was acquired on the Magpix instrument (Luminex, Austin, TX, US), and the values of analytes were calculated using ProcartaPlex Analyst 1.0 Software (ThermoFisher Scientific). The analytes with values below the lower limit of quantification in more than 75% of all samples were excluded from the analysis (**Table S1**). For the accepted analytes, the values between 0 pg/mL and the lower limit of quantification, and those exceeding the upper limit of quantification, were approximated with the lowest and highest concentration representing these limits, respectively.

#### Electrochemiluminescence assay

The following candidate biomarkers were measured using electrochemiluminescence-based human assays from Meso Scale Diagnostics (MSD, Rockville, MD, US): IL-18 (U-PLEX Human IL-18), TNF-α (U-PLEX Human TNF-α), and FGF2 (R-PLEX Human basic FGF [FGF2] Antibody Set, F210E-3). All assays were conducted according to the manufacturer’s protocols and signal detection was performed on the QuickPlex SQ 120 device (MSD). Pre-analytical data processing was done using MSD Discovery Workbench software (v4.0.12).

#### ELISA assay

The S100B ELISA Kit (#EZHS100B-33K; EMD Millipore, Billerica, MA, US) and Human MMP-14 ELISA Kit (#ab197747; Abcam, Cambridge, UK) were used to measure S100B and MMP-14 protein levels in the plasma of VS patients and controls. ELISA assays were performed by adhering to the manufacturer’s protocol. Absorbance was measured using Spectra MAX 190 plate Reader (Molecular Devices, Sunnyvale, CA, US), and the standard curve was plotted using SoftMax Pro software (v5.2; Molecular Devices, San Jose, CA, US).

### Statistics

Patient group comparisons comprised: 1) all VS patients vs. controls), 2) VS-GH patients vs. controls, 3) VS-PH patients vs. controls, and 4) VS-PH vs. VS-GH patients. For all comparisons, *P*<0.05 was considered statistically significant. Differences in age across groups were analyzed using one-way ANOVA with a Dunn’s multiple comparisons test. Differences in tumor volume and PTA were analyzed using Mann-Whitney *t* test. Differences in WR were analyzed using N-1 chi-squared test for the comparison of two proportions expressed as a percentage. The ratio method was used to assess the individual effect of candidate biomarkers on the WR of VS patients.

#### Generalized linear mixed effects regression

Generalized linear mixed effects regression models were used to assess the relationship between candidate biomarkers’ levels in the plasma of VS patients and controls vs. clinical variables. All candidate biomarkers were natural log transformed prior to modeling. To compare plasma biomarker levels in VS patients with controls while controlling for subjects’ age and sex, the following generalized least squares (GLS) model was used: ln<biomarker> =*b0* + *b1Sex* + *b2*Age + *b3*GH + *b4*PH. The biomarker in the formula refers to tested immune-related factors, while b0, b1, b2, b3, and b4 are regression coefficients representing the change in natural log-transformed concentrations of prospective biomarkers to a one-unit change in the respective independent variable (sex, age, GH, and PH). The effect of sex × candidate biomarkers interactions was assessed by the same model with the following terms: ln<biomarker> = *b0* + *b1*Sex + *b2*Age + *b3*GH + *b4*PH + *b5*Sex*ln<biomarker>. For mutual comparison of GH and PH groups, the GLS model was extended for controlling the subject’s tumor volume.

RLMs were used to regress ipsilateral PTA values on plasma biomarker levels while controlling for sex, age, tumor volume, and PTA contralateral to VS. RLMs were also used to regress tumor volume on plasma biomarker levels while controlling for sex and age. The use of RLMs was necessary for both outcomes, as non-robust linear models suffered from substantial effect size biased caused by residual outliers. RLM was used to control for patients’ age, sex, tumor size, and hearing in the contralateral ear. The model had the following terms: PTA_ipsi = *b0* + *b1*Sex + *b2*Age + *b3*PTA_contra + *b4*Tumor volume^(1/3) + *b5*ln<biomarker> where PTA_ipsi is PTA in the ear ipsilateral to the tumor and PTA_contra is PTA in the ear contralateral to the tumor. To investigate the effect of sex × candidate biomarker interactions, the same robust linear model was used with the following terms: PTA_ipsi = *b0* + *b1*Sex + *b2*Age + *b3*PTA_contra + *b4*Tumor volume^(1/3) + *b5*ln<biomarker> + *b6*Sex*ln<biomarker>. The regression coefficients (b0, b1, b2, b3, b4, b5 and b6) representing the change in PTA_ipsi to a one-unit change in the respective independent variable (sex, age, PTA_contra, tumor volume, biomarker, and interactions of sex and biomarker concentration).

In the analysis of the interaction with tumor volume, the RLM had the following terms. Tumor volume^(1/3) =*b0* + *b1*Sex + *b2*Age + *b3*ln<biomarker> where b0, b1, b2, and b3 are regression coefficients representing the change in tumor volume to a one-unit change in the respective independent variable (sex, age, and biomarker concentration). To investigate the effect of sex × candidate biomarkers interactions, the same robust linear model was extended as follows: Tumor volume^(1/3) = *b0* + *b1*Sex + *b2*Age + *b3*ln<biomarker> + *b4*Sex*<ln biomarker> where b4 is a regression coefficient representing the change in tumor volume to a one-unit change in the variable determined by interactions of sex and biomarker concentration.

FLMs were used to regress WR scores ipsilateral to VS on plasma biomarker levels while controlling for sex, age, tumor volume, and WR scores contralateral to VS. Fractional models were necessary since WR scores were measured on the interval [0, 100] and therefore had hard upper and lower bounds, as well as being heteroskedastic. FLM was used to control for patients’ age, sex, tumor size, and hearing in the contralateral ear. The model had the following terms: WR_ipsi = *b0* + *b1*Sex + *b2*Age + *b3*WR_contra + *b4*Tumor volume^(1/3) *+ b5*ln<biomarker> where WR_ipsi is WR in the ear ipsilateral to the tumor and WR_contra is WR in the ear contralateral to the tumor. The effect of sex × candidate biomarkers interactions was assessed by the same model with the following terms: WR_ipsi = *b0* + *b1*Sex + *b2*Age + *b3*WR_contra + *b4*Tumor volume^(1/3) *+ b5*ln<biomarker> + *b6*Sex*ln<biomarker>. The regression coefficients (b0, b1, b2, b3, b4, b5 and b6) represent the change in WR_ipsi to a one-unit change in the respective independent variable (sex, age, WR_contra, tumor volume, biomarker, and interactions of sex and biomarker concentration).

ROC analysis was used to evaluate the diagnostic power and utility of significantly elevated candidate plasma biomarkers. The ROC curves were constructed and the areas under the ROC curve (AUC) were calculated using MedCalc software. The discriminatory power of candidate biomarkers was categorized as follows: “outstanding” discrimination, AUC ≥ 0.90; “excellent” discrimination, 0.80 ≤ AUC < 0.90; “acceptable” discrimination, 0.70 ≤ AUC < 0.80; and “poor” discrimination, <0.70 (*56*). The candidate biomarkers with a similar association with hearing parameters, tumor size, and similar degree of elevated plasma levels were compared mutually by precision-recall curves analysis using MedCalc software.

The predictability of biomarkers was verified by precision-recall curve analysis, where the individual curves were compared between biomarkers belonging to the same category based on their association with hearing, tumor size, and the extent of elevation in VS patients.

#### Statistical software

Comparisons of demographics, tumor volume, and hearing loss between groups were performed with GraphPad Prism (v9.3.1; GraphPad Software, La Jolla, CA, US) and MedCalc^®^ Statistical Software (v20.109; MedCalc Software Ltd, Ostend, Belgium). Models were estimated and graphs were generated using R (v3.6.3; R Foundation, Vienna, Austria). Combinatorial analysis of candidate biomarkers was performed using the CombiROC web application (http://combiroc.eu) (*57*) and verified by logistic regression with the enter method using MedCalc® software. The relationship between panel biomarkers was assessed by Spearman correlation analysis using GraphPad software. Protein fold change was calculated in Microsoft Excel 2013 as described by Augulian et al. (*58*), using the Real Statistics Resource Pack plugin for Excel developed by Dr. Charles Zaiont and available at http://www.real-statistics.com.

### Study approval

All study protocols were approved by the Human Studies Committee of MEEI and MGH (IRB protocol #14-148H). All participants provided written informed consent prior to participation.

## Supporting information

Supplementary_materials_Vasilijic_et_al_2022_v1

## Acknowledgments

We thank Shelley Batts, PhD of Stanford University for assistance with scientific writing.

## Author contributions

Conceptualization: KMS Sample providing: RL, DBW; Sample processing: SV, NA, HH, YR, JS, MS, TF, LL; Creation of the VS patient database: SV, NA, HH, YR, JS, MS, TF, LL; MRI analyses: HH; Data analysis: SV, SW, KMS; Visualization: SV; Supervision: KMS; Writing—original draft: SV, KMS; Writing— review & editing: All authors

## Data and materials availability

All data are available in the main text or the supplementary materials.

