## Supplementary_materials_Vasilijic_et_al_2022_v1 for "Identification of Immune-Related Candidate Biomarkers in Plasma of Patients with Sporadic Vestibular Schwannoma"

Sasa Vasilijic *et al.*

**This PDF file includes:**

Figs. S1 to S10
Tables S1 to S4

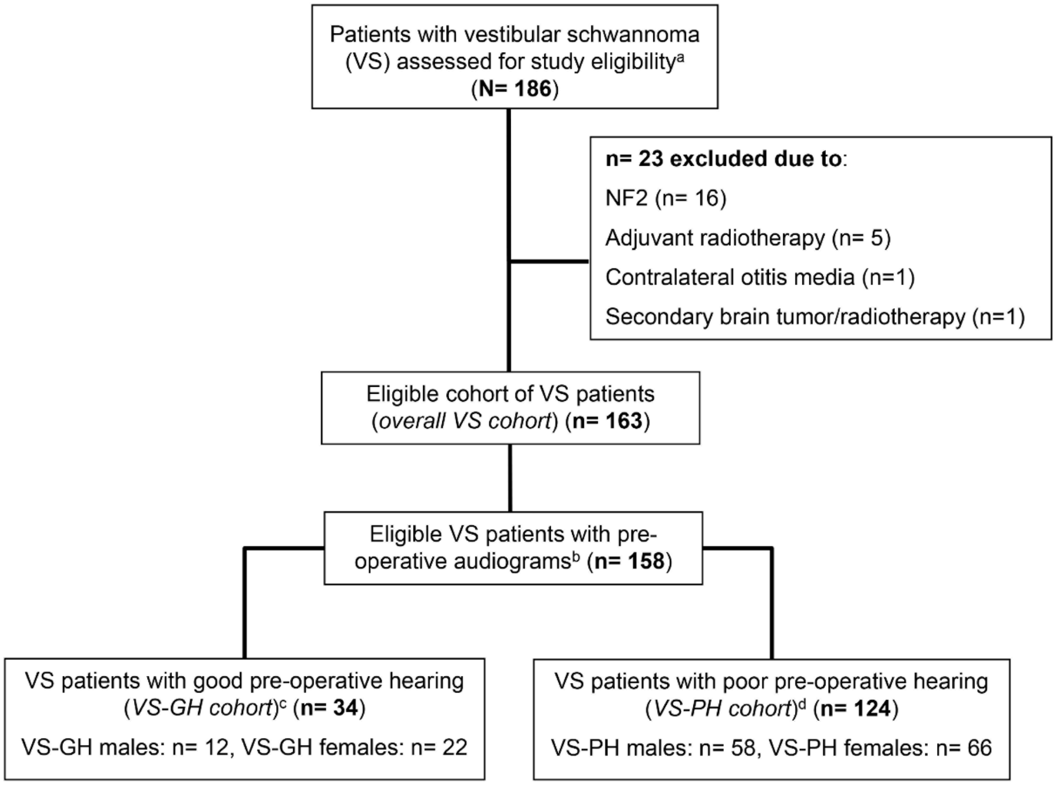


**Fig. S1. CONSORT flow diagram for the VS cohort**

Sample selection flowchart for the VS cohort. Among the overall cohort of eligible VS patients, two subgroups were further defined by pre-operative hearing ability.

Notes: [a] Eligible patients had unilateral, sporadic VS that had not been previously resected or irradiated; [b] Five VS-PH patients had no pre-operative audiograms. [c] GH was defined as word recognition score >70% and pure tone average <30 decibels (dB). [d] PH was defined as word recognition score ≤70% and pure tone average ≥30 dB. Abbreviation: NF2, neurofibromatosis 2.

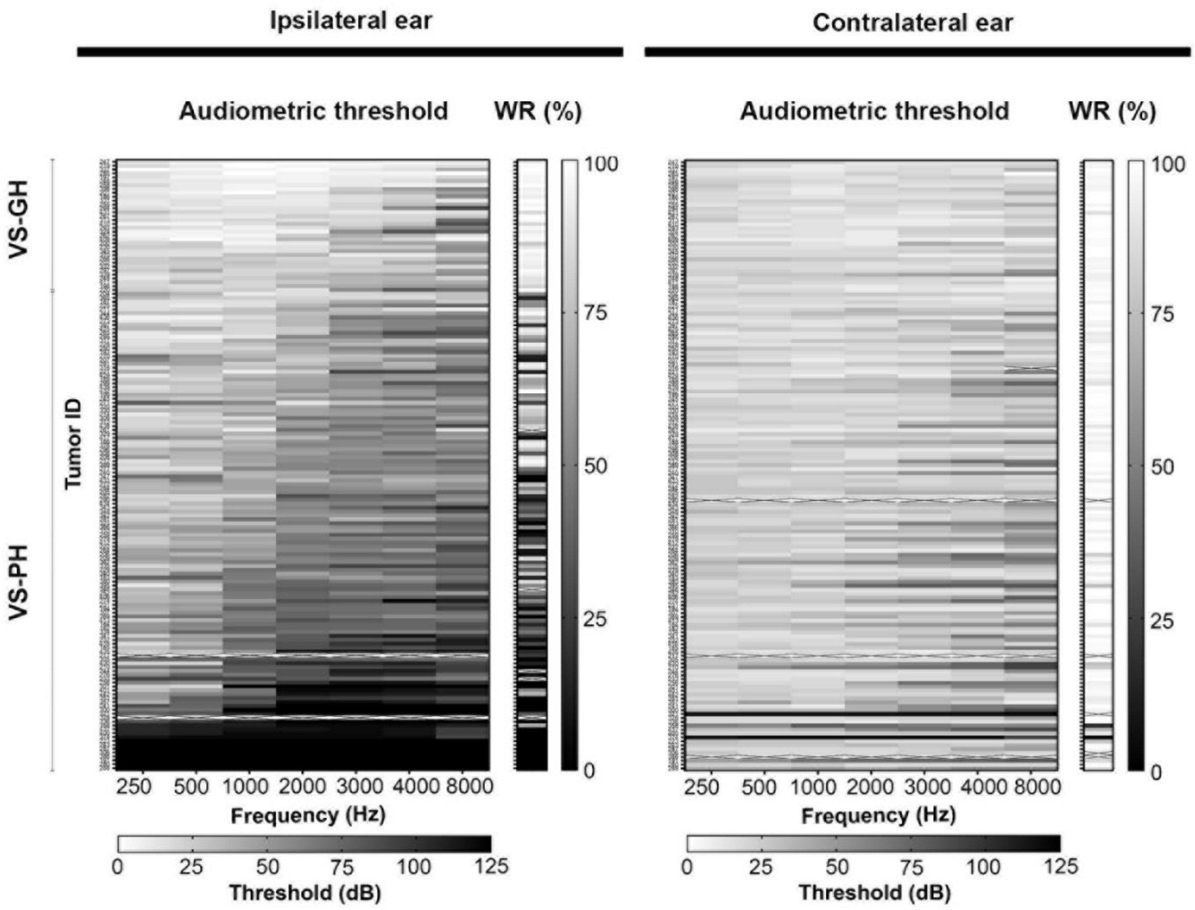


**Fig. S2. Hearing characteristics of VS patients**

Abbreviation: dB, decibel; GH, good hearing; Hz, hertz; PH, poor hearing; VS, vestibular schwannoma; WR, word recognition.

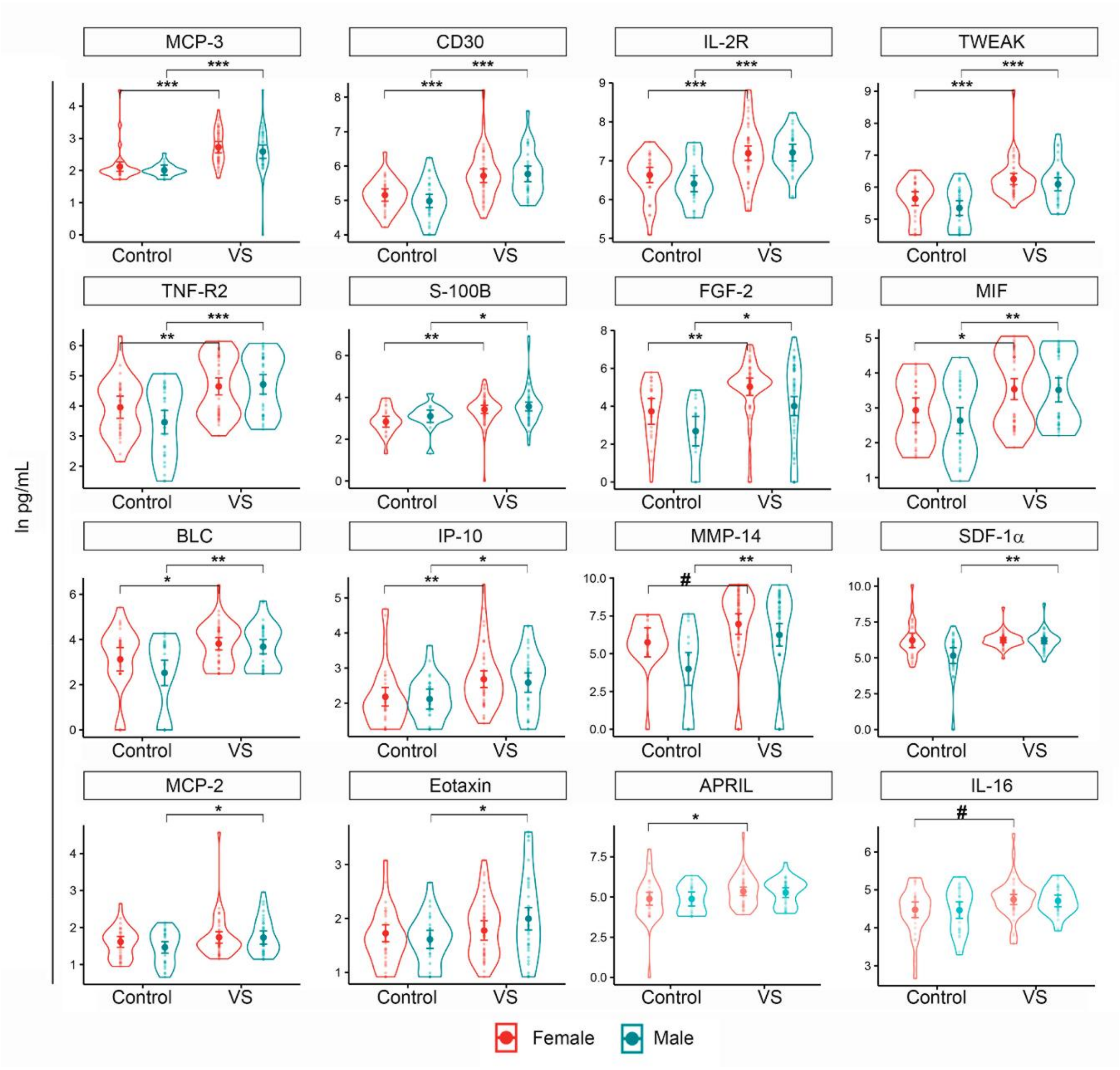


**Fig. S3. Significantly elevated candidate biomarkers in female and male VS patients**

Abbreviations: VS, vestibular schwannoma. **Padj*<0.05, ***Padj*<0.01, ****Padj*<0.001, #significant at *P*<0.05 prior to adjustment.

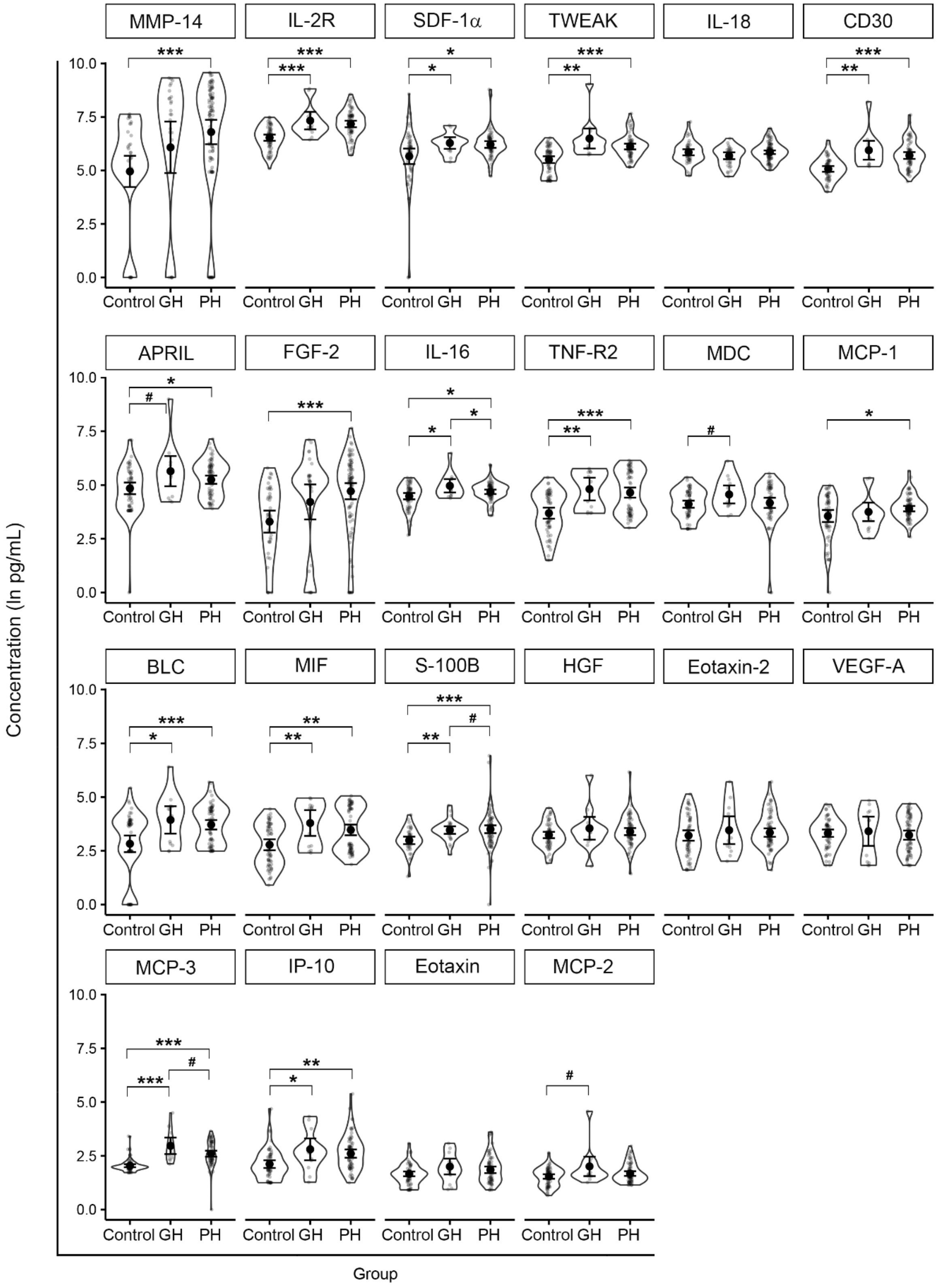

MMP-14
IL-2R
SDF-1α
TWEAK
IL-18
CD30
APRIL
FGF-2
IL-16
TNF-R2
MDC
MCP-1
BLC
MIF
S-100B
HGF
Eotaxin-2
VEGF-A
MCP-3
IP-10
Eotaxin
MCP-2
Concentration (ln pg/mL)
Control GH PH
Group

**Fig. S4. Plasma levels of candidate biomarkers in VS-GH patients, VS-PH patients, and controls**

Abbreviations: GH, good hearing; PH, poor hearing; VS, vestibular schwannoma. *$P_{adj}$<0.05, **$P_{adj}$<0.01, ***$P_{adj}$<0.001, #significant at $P$<0.05 prior to adjustment.

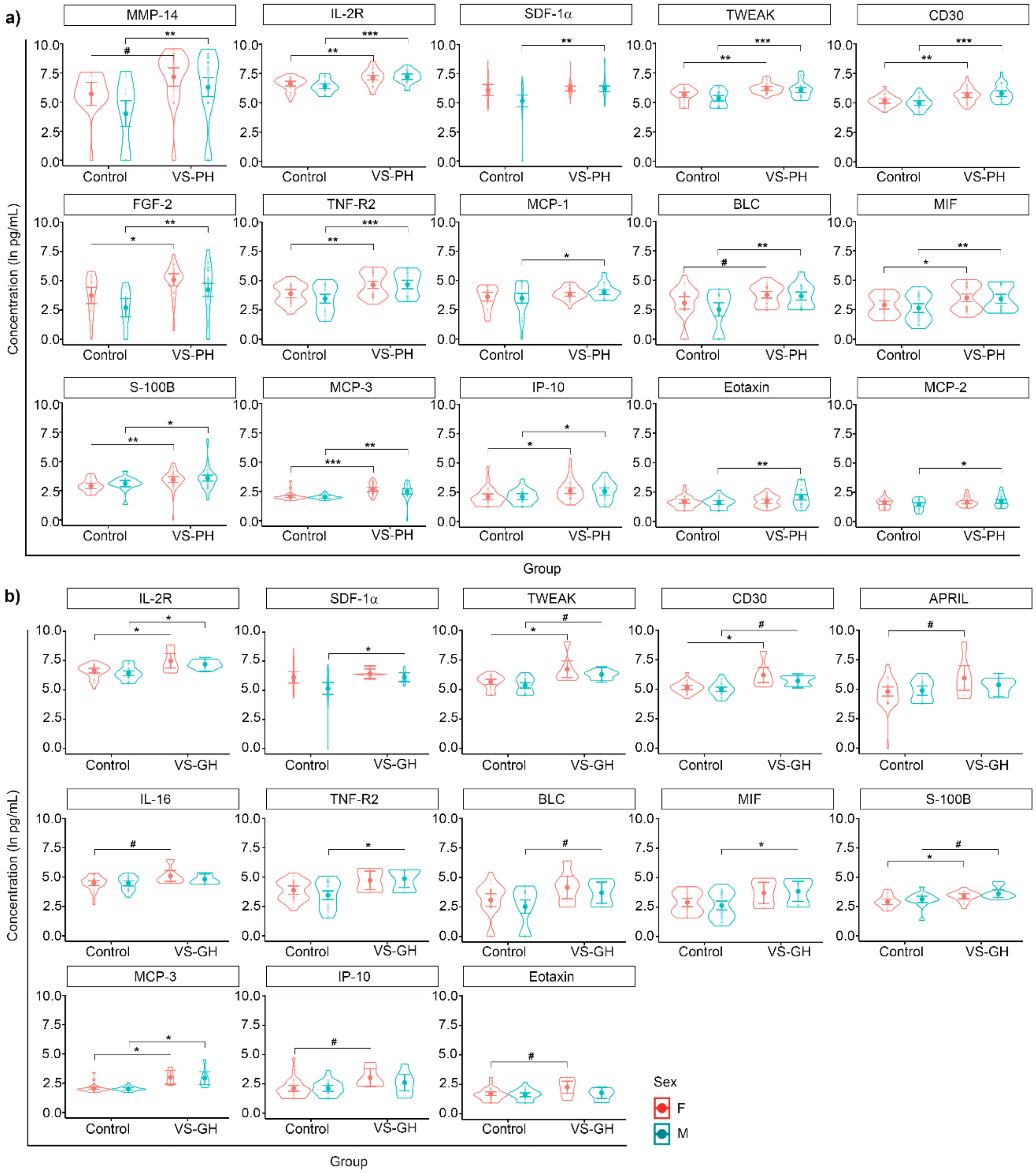


**Fig. S5. Significantly elevated candidate biomarkers in female and male VS patients with (a) poor or (b) good hearing compared to controls**

Abbreviations: GH, good hearing; PH, poor hearing; VS, vestibular schwannoma. *$Padj<0.05$, **$Padj<0.01$, ***$Padj<0.001$, #significant at $P<0.05$ prior to adjustment.

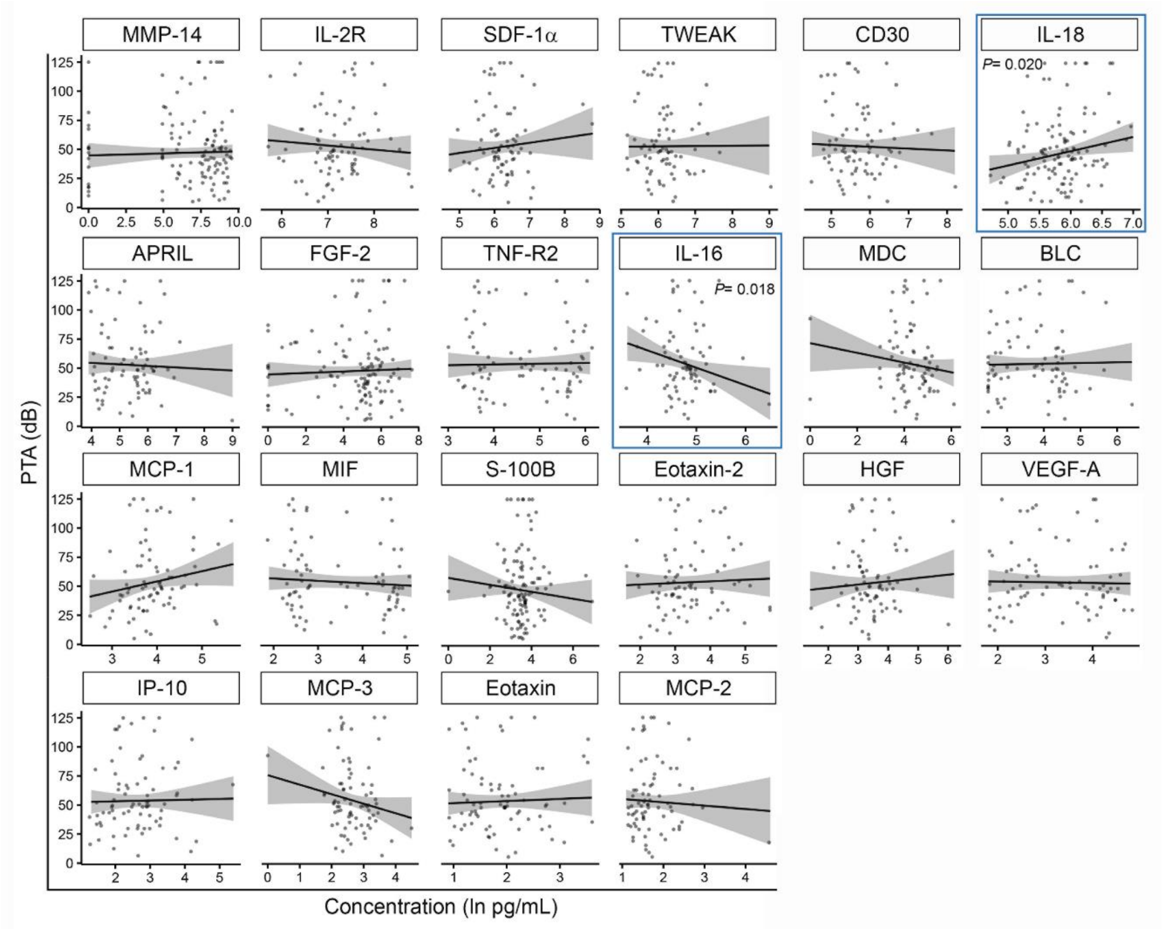


**Fig. S6. Plasma level of candidate biomarkers and severity of pre-operative pure tone average in VS patients**

The blue box indicates significant associations for IL-18 and IL-16 prior to p-value adjustment. Abbreviations: PTA, pure-tone average; VS, vestibular schwannoma.

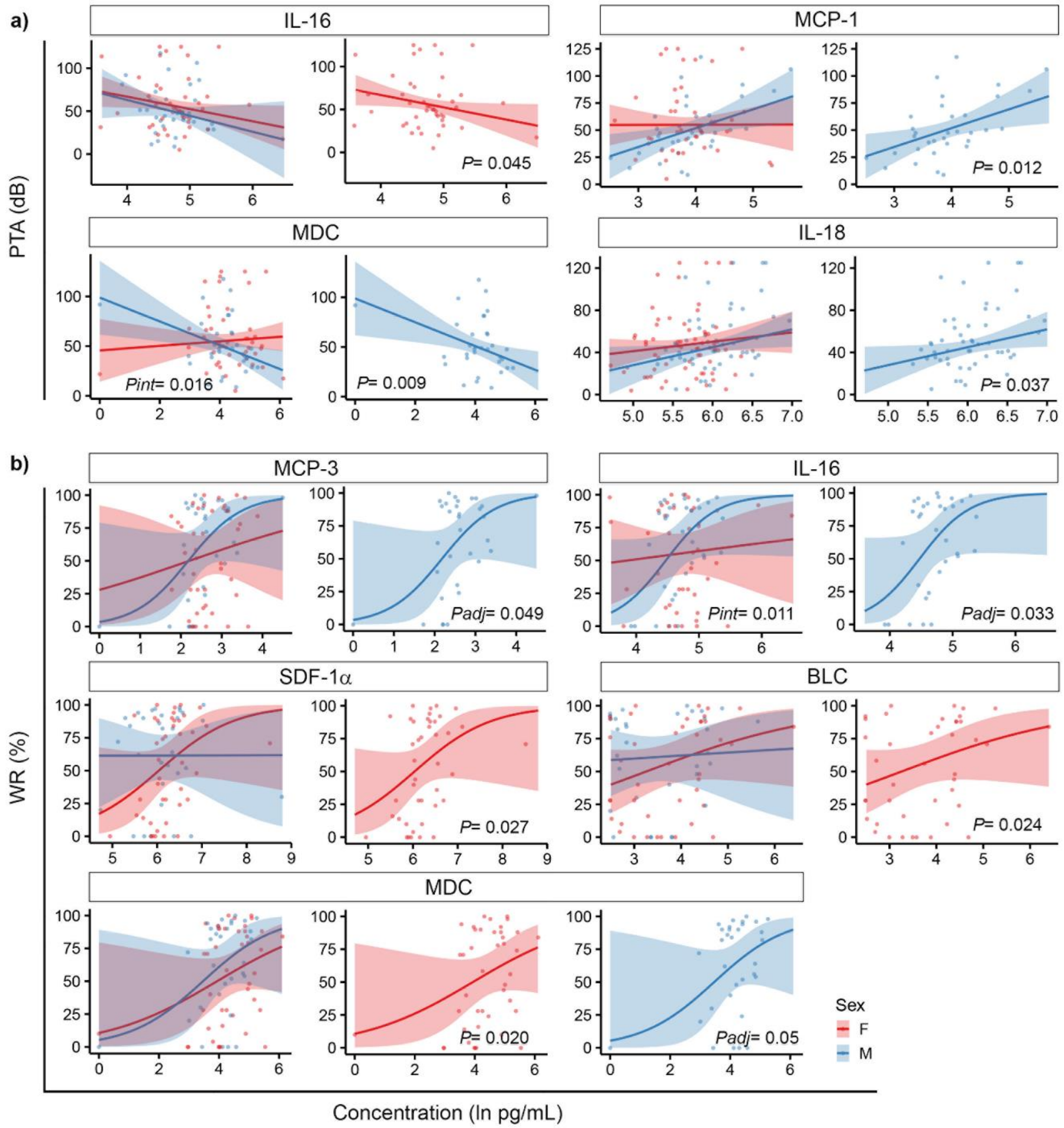


**Fig. S7. Sex differences in the association of plasma biomarker levels and severity of pre-operative a) PTA and b) WR in VS patients**

Abbreviations: dB, decibels; PTA, pure-tone average; VS, vestibular schwannoma; WR, word recognition.

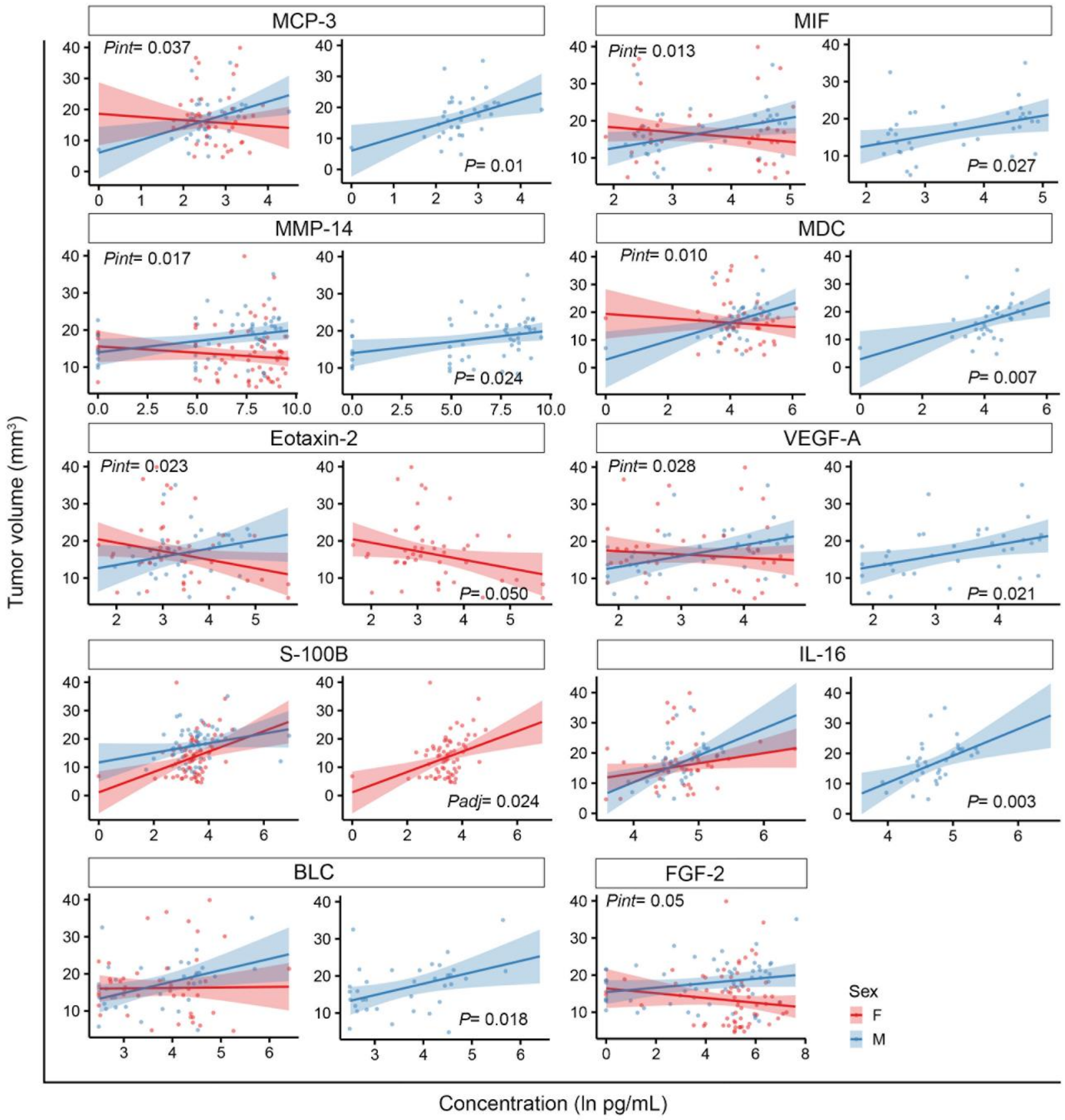


**Fig. S8. Sex differences in the association of plasma biomarker levels and tumor volume in VS patients**

Abbreviation: VS, vestibular schwannoma.

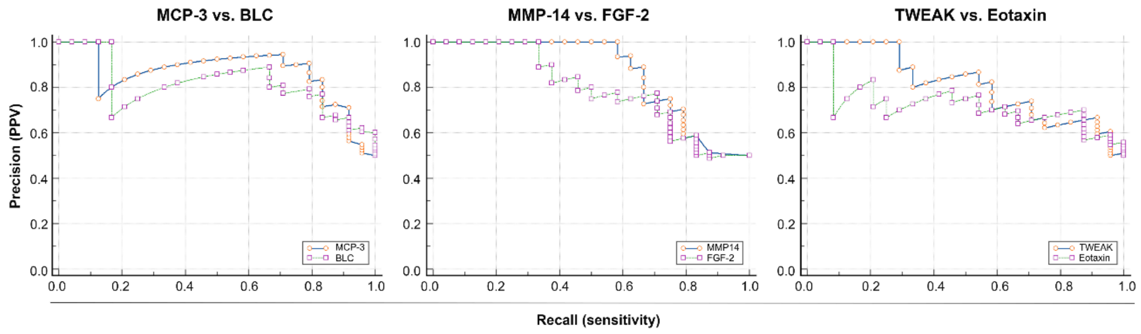


**Fig. S9. Precision-recall analysis**

Abbreviation: PPV, positive predictive value.

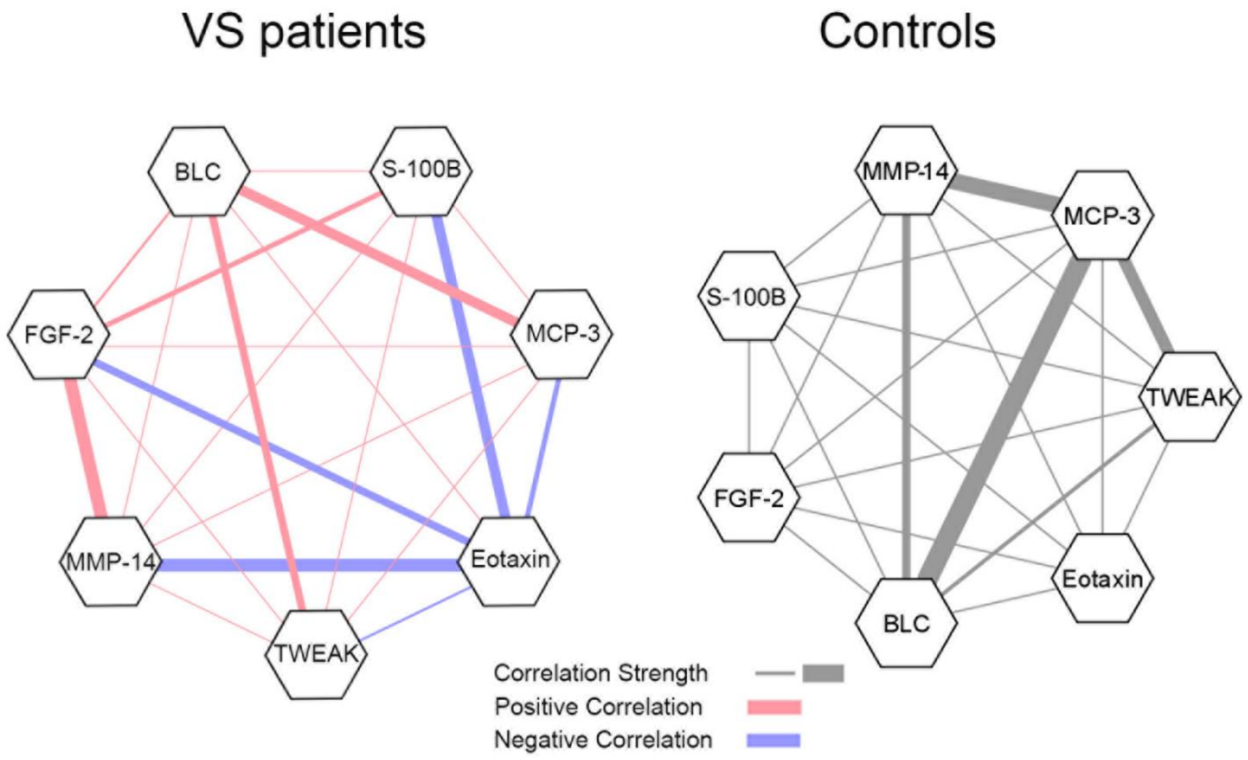


**Fig. S10. Correlation networks of candidate biomarkers**

Abbreviation: VS, vestibular schwannoma.

**Table S1. List and full names of assayed immune factors**

| Common name | Full name | Common name | Full name | Common name | Full name |
|---|---|---|---|---|---|
| APRIL | A proliferation-inducing ligand | IL-1α | Interleukin-1α | MDC | Macrophage-derived chemokine (CCL22) |
| BAFF | B-cell activating factor | IL-1β | Interleukin-1β | MIF | Macrophage migration inhibitory factor |
| BLC | B lymphocyte chemoattractant (CXCL13) | IL-2 | Interleukin-2 | MIG | Monokine induced by interferon gamma (CXCL9) |
| CD30 | CD30 | IL-20 | Interleukin-20 | MIP-1α | Macrophage inflammatory protein-1α (CCL3) |
| CD40L | CD40 ligand | IL-21 | Interleukin-21 | MIP-1β | Macrophage inflammatory protein-1β (CCL4) |
| ENA-78 | Epithelial neutrophil-activating peptide 78 (CXCL5) | IL-22 | Interleukin-22 | MIP-3α | Macrophage inflammatory protein-3α (CCL20) |
| Eotaxin | Eotaxin (CCL11) | IL-23 | Interleukin-23 | MMP-1 | Matrix metalloproteinase-1 |
| Eotaxin-2 | Eotaxin-2 (CCL24) | IL-27 | Interleukin-27 | MMP-14 | Matrix metalloproteinase-14 |
| Eotaxin-3 | Eotaxin-3 (CCL26) | IL-2R | Interleukin-2R | NGF- β | Nerve Growth Factor-β |
| FGF-2 | Fibroblast Growth Factor-2 | IL-3 | Interleukin-3 | S100B | S100B |
| Fractalkine | Fractalkine (CX3CL1) | IL-31 | Interleukin-31 | SCF | Stem cell factor |
| G-CSF | Granulocyte-colony stimulating factor | IL-4 | Interleukin-4 | SDF-1α | Stromal cell-derived factor 1α (CXCL12) |
| GM-CSF | Granulocyte macrophage colony-stimulating factor | IL-5 | Interleukin-5 | TNF-α | Tumor necrosis factor- α |
| GRO-α | Growth-related oncogene α (CXCL1) | IL-6 | Interleukin-6 | TNF- β | Tumor necrosis factor- β |
| HGF | Hepatocyte Growth Factor | IL-7 | Interleukin-7 | TNF-R2 | Tumor necrosis factor- receptor 2 |
| IFN-α | Interferon-α | IL-8 | Interleukin-8 | TRAIL | Tumor necrosis factor (TNF)-related apoptosis-inducing ligand |
| IFN-g | Interferon-ɣ | IL-9 | Interleukin-9 | TSLP | Thymic stromal lymphopoietin |
| IL-10 | Interleukin-10 | IP-10 | Interferon-γ inducible protein-10 (CXCL10) | TWEAK | Tumor necrosis factor-like weak inducer of apoptosis |
| IL-12p70 | Interleukin-12p70 | I-TAC | Interferon–inducible T Cell Alpha | VEGF-A | Vascular endothelial growth factor-A |

| | | | |
|---|---|---|---|
| | | | Chemoattractant (CXCL11) |
| IL-13 | Interleukin-13 | LIF | Leukemia inhibitory factor |
| IL-15 | Interleukin-15 | MCP-1 | Monocyte chemoattractant protein-1 (CCL2) |
| IL-16 | Interleukin-16 | MCP-2 | Monocyte chemoattractant protein-2 (CCL8) |
| IL-17A | Interleukin-17A | MCP-3 | Monocyte chemoattractant protein-3 (CCL7) |
| IL-18 | Interleukin-18 | M-CSF | Macrophage colony-stimulating factor |

**Table S2. Detectability of candidate biomarkers in plasma of VS patients**

| Candidate biomarkers | Samples between LLOQ and ULOQ | Candidate biomarkers | Samples between LLOQ and ULOQ | Candidate biomarkers | Samples between LLOQ and ULOQ |
|---|---|---|---|---|---|
| **APRIL** | **81/81 (100%)** [a] | IL-18 | 43/81 (53%) [a] | **MCP-3** | **79/81 (98%)** [a] |
| BAFF | 56/81 (69%) [a] | **IL-18** | **123/123 (100%)** [b] | M-CSF | 8/81 (10%) [a] |
| **BLC** | **73/81 (90%)** [a] | IL-1α | 8/81 (10%) [a] | **MDC** | **77/81 (95%)** [a] |
| **CD30** | **81/81 (100%)** [a] | IL-1β | 6/81 (7%) [a] | **MIF** | **81/81 (100%)** [a] |
| CD40L | 32/81(40%) [a] | IL-2 | 49/81 (60%) [a] | MIG | 30/81 (37%) [a] |
| ENA-78 | 49/81 (60%) [a] | IL-20 | 19/81 (23%) [a] | MIP-1α | 43/81 (53%) [a] |
| **Eotaxin** | **77/81 (95%)** [a] | IL-21 | 7/81 (9%) [a] | MIP-1β | 36/81 (44%) [a] |
| **Eotaxin-2** | **80/81 (99%)** [a] | IL-22 | 6/81 (7%) [a] | MIP-3α | 5/81 (6%) [a] |
| Eotaxin-3 | 47/81 (58%) [a] | IL-23 | 14/81 (17%) [a] | MMP1 | 7/81 (9%) [a] |
| FGF-2 | 35/81 (43%) [a] | IL-27 | 16/81 (20%) [a] | **MMP-14** | **97/123 (79%)** [c] |
| **FGF-2** | **111/123 (90%)** [b] | **IL-2R** | **81/81 (100%)** [a] | NGF- β | 10/81 (12%) [a] |
| Fractalkine | 10/81 (12%) [a] | IL-3 | 4/81 (5%) [a] | **S100 B** | **122/123 (99%)** [c] |
| G-CSF | 6/81 (7%) [a] | IL-31 | 5/81 (6%) [a] | SCF | 19/81 (23%) [a] |
| GM-CSF | 2/81 (2%) [a] | IL-4 | 15/81 (19%) [a] | **SDF-1α** | **81/81 (100%)** [a] |
| GRO-α | 1/81 (1%) [a] | IL-5 | 7/81 (9%) [a] | TNF-α | 3/81 (4%) [a] |
| **HGF** | **80/81 (99%)** [a] | IL-6 | 8/81 (10%) [a] | TNF-α | 29/123 (24%) [b] |
| IFN-α | 5/81 (6%) [a] | IL-7 | 14/81 (17%) [a] | TNF- β | 14/81 (17%) [a] |
| IFN-γ | 21/81 (26%) [a] | IL-8 | 15/81 (19%) [a] | **TNF-R2** | **81/81 (100%)** [a] |
| IL-10 | 16/81 (20%) [a] | IL-9 | 5/81 (6%) [a] | TRAIL | 49/81 (60%) [a] |
| IL-12p70 | 4/81 (5%) [a] | **IP-10** | **80/81 (99%)** [a] | TSLP | 15/81 (19%) [a] |
| IL-13 | 18/81 (22%) [a] | I-TAC | 5/81 (6%) [a] | **TWEAK** | **81/81 (100%)** [a] |
| IL-15 | 13/81 (16%) [a] | LIF | 18/81 (22%) [a] | **VEGF-A** | **76/81 (94%)** [a] |
| **IL-16** | **81/81 (100%)** [a] | **MCP-1** | **81/81 (100%)** [a] | | |
| IL-17A | 33/81 (41%) [a] | **MCP-2** | **80/81 (99%)** [a] | | |

Candidate biomarkers with an absolute concentration between the LLOQ and ULOQ calculated in 75% or more of tested plasma samples were included in the study. Twenty-two of 67 tested candidate biomarkers fulfilled the inclusion criteria (bold). Included candidate biomarkers whose absolute concentration was between 0 pg/mL and LLOQ were assigned with the value of the corresponding LLOQ. Candidate biomarkers exceeding ULOQ were approximated with the highest concentration representing these limits. Abbreviations: LLOQ, lower limit of quantification; ULOQ, upper limit of quantification; VS, vestibular schwannoma. Notes: [a] Assessed with Luminex assay; [b] Assessed with electrochemiluminescence assay; [c] Assessed with ELISA assay.

### Table S3. Combinatorial analysis (CombiROC) of seven selected candidate biomarkers

| Biomarker combinations | Symbol | AUC | SE | SP | Opt Cutoff |
|---|---|---|---|---|---|
| MCP3 | Marker_1 | 0.862 | 0.792 | 0.917 | 0.554 |
| BLC | Marker_2 | 0.836 | 0.833 | 0.75 | 0.34 |
| S100B | Marker_3 | 0.729 | 0.625 | 0.792 | 0.538 |
| FGF2 | Marker_4 | 0.731 | 0.708 | 0.792 | 0.563 |
| MMP14 | Marker_5 | 0.792 | 0.625 | 0.958 | 0.607 |
| TWEAK | Marker_6 | 0.786 | 0.708 | 0.75 | 0.525 |
| Eotaxin | Marker_7 | 0.752 | 0.875 | 0.625 | 0.414 |
| MCP3-BLC | Combo I | 0.861 | 0.75 | 0.958 | 0.7 |
| MCP3-S100B | Combo II | 0.88 | 0.917 | 0.833 | 0.282 |
| MCP3-FGF2 | Combo III | 0.862 | 0.833 | 0.875 | 0.401 |
| MCP3-MMP14 | Combo IV | 0.879 | 0.875 | 0.875 | 0.417 |
| MCP3-TWEAK | Combo V | 0.873 | 0.833 | 0.875 | 0.462 |
| MCP3-Eotaxin | Combo VI | 0.913 | 0.875 | 0.875 | 0.411 |
| BLC-S100B | Combo VII | 0.825 | 0.875 | 0.792 | 0.404 |
| BLC-FGF2 | Combo VIII | 0.835 | 0.833 | 0.75 | 0.359 |
| BLC-MMP14 | Combo IX | 0.828 | 0.75 | 0.833 | 0.52 |
| BLC-TWEAK | Combo X | 0.849 | 0.792 | 0.792 | 0.377 |
| BLC-Eotaxin | Combo XI | 0.88 | 0.958 | 0.75 | 0.297 |
| S100B-FGF2 | Combo XII | 0.773 | 0.792 | 0.708 | 0.437 |
| S100B-MMP14 | Combo XIII | 0.757 | 0.792 | 0.75 | 0.449 |
| S100B-TWEAK | Combo XIV | 0.823 | 0.958 | 0.583 | 0.279 |
| S100B-Eotaxin | Combo XV | 0.823 | 0.875 | 0.708 | 0.422 |
| FGF2-MMP14 | Combo XVI | 0.775 | 0.542 | 1 | 0.626 |
| FGF2-TWEAK | Combo XVII | 0.79 | 0.833 | 0.708 | 0.44 |
| FGF2-Eotaxin | Combo XVIII | 0.826 | 0.875 | 0.667 | 0.475 |
| MMP14-TWEAK | Combo XIX | 0.807 | 0.625 | 0.917 | 0.599 |
| MMP14-Eotaxin | Combo XX | 0.812 | 0.958 | 0.625 | 0.317 |
| TWEAK-Eotaxin | Combo XXI | 0.854 | 0.833 | 0.792 | 0.501 |
| MCP3-BLC-S100B | Combo XXII | 0.882 | 0.917 | 0.833 | 0.31 |
| MCP3-BLC-FGF2 | Combo XXIII | 0.861 | 0.833 | 0.875 | 0.404 |
| MCP3-BLC-MMP14 | Combo XXIV | 0.878 | 0.875 | 0.875 | 0.418 |
| MCP3-BLC-TWEAK | Combo XXV | 0.882 | 0.833 | 0.958 | 0.551 |
| **MCP3-BLC-Eotaxin** | **Combo XXVI** | **0.913** | **0.875** | **0.875** | **0.407** |
| MCP3-S100B-FGF2 | Combo XXVII | 0.884 | 0.917 | 0.833 | 0.266 |

| | | | | | |
|---|---|---|---|---|---|
| **MCP3-S100B-MMP14** | **Combo XXVIII** | **0.901** | **0.792** | **0.917** | **0.617** |
| MCP3-S100B-TWEAK | Combo XXIX | 0.884 | 0.792 | 0.917 | 0.553 |
| **MCP3-S100B-Eotaxin** | **Combo XXX** | **0.918** | **1** | **0.708** | **0.183** |
| MCP3-FGF2-MMP14 | Combo XXXI | 0.876 | 0.875 | 0.875 | 0.459 |
| MCP3-FGF2-TWEAK | Combo XXXII | 0.873 | 0.833 | 0.875 | 0.465 |
| **MCP3-FGF2-Eotaxin** | **Combo XXXIII** | **0.906** | **0.792** | **0.958** | **0.721** |
| MCP3-MMP14-TWEAK | Combo XXXIV | 0.887 | 0.792 | 0.958 | 0.713 |
| **MCP3-MMP14-Eotaxin** | **Combo XXXV** | **0.908** | **0.833** | **0.958** | **0.638** |
| **MCP3-TWEAK-Eotaxin** | **Combo XXXVI** | **0.91** | **0.875** | **0.875** | **0.469** |
| BLC-S100B-FGF2 | Combo XXXVII | 0.823 | 0.875 | 0.792 | 0.408 |
| BLC-S100B-MMP14 | Combo XXXVIII | 0.844 | 0.875 | 0.792 | 0.385 |
| BLC-S100B-TWEAK | Combo XXXIX | 0.861 | 0.875 | 0.75 | 0.354 |
| BLC-S100B-Eotaxin | Combo XL | 0.88 | 1 | 0.75 | 0.286 |
| BLC-FGF2-MMP14 | Combo XLI | 0.833 | 0.792 | 0.833 | 0.484 |
| BLC-FGF2-TWEAK | Combo XLII | 0.842 | 0.75 | 0.833 | 0.444 |
| BLC-FGF2-Eotaxin | Combo XLIII | 0.882 | 0.958 | 0.75 | 0.301 |
| BLC-MMP14-TWEAK | Combo XLIV | 0.839 | 0.75 | 0.833 | 0.439 |
| BLC-MMP14-Eotaxin | Combo XLV | 0.877 | 0.958 | 0.75 | 0.305 |
| BLC-TWEAK-Eotaxin | Combo XLVI | 0.88 | 0.917 | 0.792 | 0.355 |
| S100B-FGF2-MMP14 | Combo XLVII | 0.774 | 0.833 | 0.708 | 0.404 |
| S100B-FGF2-TWEAK | Combo XLVIII | 0.802 | 0.833 | 0.75 | 0.397 |
| S100B-FGF2-Eotaxin | Combo XLIX | 0.844 | 0.75 | 0.833 | 0.622 |
| S100B-MMP14-TWEAK | Combo L | 0.83 | 0.75 | 0.875 | 0.532 |
| S100B-MMP14-Eotaxin | Combo LI | 0.845 | 0.958 | 0.667 | 0.272 |
| S100B-TWEAK-Eotaxin | Combo LII | 0.865 | 0.792 | 0.833 | 0.536 |
| FGF2-MMP14-TWEAK | Combo LIII | 0.806 | 0.75 | 0.792 | 0.458 |
| FGF2-MMP14-Eotaxin | Combo LIV | 0.826 | 0.833 | 0.75 | 0.518 |
| FGF2-TWEAK-Eotaxin | Combo LV | 0.865 | 0.75 | 0.833 | 0.593 |
| MMP14-TWEAK-Eotaxin | Combo LVI | 0.866 | 0.75 | 0.833 | 0.494 |
| MCP3-BLC-S100B-FGF2 | Combo LVII | 0.885 | 0.917 | 0.833 | 0.305 |
| **MCP3-BLC-S100B-MMP14** | **Combo LVIII** | **0.901** | **0.917** | **0.792** | **0.294** |
| **MCP3-BLC-S100B-TWEAK** | **Combo LIX** | **0.908** | **0.833** | **0.958** | **0.52** |
| **MCP3-BLC-S100B-Eotaxin** | **Combo LX** | **0.92** | **0.958** | **0.792** | **0.314** |
| MCP3-BLC-FGF2-MMP14 | Combo LXI | 0.878 | 0.875 | 0.917 | 0.476 |
| MCP3-BLC-FGF2-TWEAK | Combo LXII | 0.882 | 0.833 | 0.958 | 0.532 |
| **MCP3-BLC-FGF2-Eotaxin** | **Combo LXIII** | **0.901** | **0.792** | **0.958** | **0.735** |
| MCP3-BLC-MMP14-TWEAK | Combo LXIV | 0.889 | 0.833 | 0.958 | 0.625 |

| | | | | | |
|---|---|---|---|---|---|
| MCP3-BLC-MMP14-Eotaxin | Combo LXV | 0.906 | 0.875 | 0.917 | 0.611 |
| MCP3-BLC-TWEAK-Eotaxin | Combo LXVI | 0.901 | 0.875 | 0.917 | 0.476 |
| MCP3-S100B-FGF2-MMP14 | Combo LXVII | 0.898 | 0.792 | 0.917 | 0.604 |
| MCP3-S100B-FGF2-TWEAK | Combo LXVIII | 0.899 | 0.833 | 0.875 | 0.502 |
| **MCP3-S100B-FGF2-Eotaxin** | **Combo LXIX** | **0.917** | **0.958** | **0.75** | **0.246** |
| **MCP3-S100B-MMP14-TWEAK** | **Combo LXX** | **0.906** | **0.792** | **0.917** | **0.629** |
| **MCP3-S100B-MMP14-Eotaxin** | **Combo LXXI** | **0.92** | **0.875** | **0.917** | **0.564** |
| **MCP3-S100B-TWEAK-Eotaxin** | **Combo LXXII** | **0.924** | **1** | **0.708** | **0.193** |
| MCP3-FGF2-MMP14-TWEAK | Combo LXXIII | 0.891 | 0.875 | 0.875 | 0.473 |
| **MCP3-FGF2-MMP14-Eotaxin** | **Combo LXXIV** | **0.905** | **0.833** | **0.958** | **0.663** |
| **MCP3-FGF2-TWEAK-Eotaxin** | **Combo LXXV** | **0.908** | **0.875** | **0.875** | **0.466** |
| **MCP3-MMP14-TWEAK-Eotaxin** | **Combo LXXVI** | **0.911** | **0.833** | **0.958** | **0.652** |
| BLC-S100B-FGF2-MMP14 | Combo LXXVII | 0.839 | 0.875 | 0.792 | 0.393 |
| BLC-S100B-FGF2-TWEAK | Combo LXXVIII | 0.852 | 0.875 | 0.75 | 0.364 |
| BLC-S100B-FGF2-Eotaxin | Combo LXXIX | 0.882 | 0.958 | 0.792 | 0.309 |
| BLC-S100B-MMP14-TWEAK | Combo LXXX | 0.854 | 0.875 | 0.708 | 0.32 |
| BLC-S100B-MMP14-Eotaxin | Combo LXXXI | 0.88 | 1 | 0.75 | 0.282 |
| BLC-S100B-TWEAK-Eotaxin | Combo LXXXII | 0.887 | 0.958 | 0.75 | 0.345 |
| BLC-FGF2-MMP14-TWEAK | Combo LXXXIII | 0.842 | 0.75 | 0.833 | 0.44 |
| BLC-FGF2-MMP14-Eotaxin | Combo LXXXIV | 0.88 | 0.833 | 0.833 | 0.406 |
| BLC-FGF2-TWEAK-Eotaxin | Combo LXXXV | 0.884 | 0.833 | 0.833 | 0.5 |
| BLC-MMP14-TWEAK-Eotaxin | Combo LXXXVI | 0.892 | 0.917 | 0.792 | 0.371 |
| S100B-FGF2-MMP14-TWEAK | Combo LXXXVII | 0.83 | 0.792 | 0.833 | 0.485 |
| S100B-FGF2-MMP14-Eotaxin | Combo LXXXVIII | 0.856 | 0.833 | 0.75 | 0.479 |
| S100B-FGF2-TWEAK-Eotaxin | Combo LXXXIX | 0.877 | 0.917 | 0.708 | 0.327 |
| S100B-MMP14-TWEAK-Eotaxin | Combo XC | 0.873 | 0.875 | 0.75 | 0.405 |
| FGF2-MMP14-TWEAK-Eotaxin | Combo XCI | 0.865 | 1 | 0.583 | 0.226 |
| MCP3-BLC-S100B-FGF2-MMP14 | Combo XCII | 0.896 | 0.75 | 0.958 | 0.727 |
| **MCP3-BLC-S100B-FGF2-TWEAK** | **Combo XCIII** | **0.908** | **0.833** | **0.958** | **0.536** |
| **MCP3-BLC-S100B-FGF2-Eotaxin** | **Combo XCIV** | **0.917** | **0.958** | **0.792** | **0.306** |
| **MCP3-BLC-S100B-MMP14-TWEAK** | **Combo XCV** | **0.908** | **0.875** | **0.875** | **0.463** |
| **MCP3-BLC-S100B-MMP14-Eotaxin** | **Combo XCVI** | **0.922** | **0.875** | **0.875** | **0.478** |
| **MCP3-BLC-S100B-TWEAK-Eotaxin** | **Combo XCVII** | **0.927** | **0.875** | **0.958** | **0.557** |
| MCP3-BLC-FGF2-MMP14-TWEAK | Combo XCVIII | 0.892 | 0.875 | 0.958 | 0.533 |
| **MCP3-BLC-FGF2-MMP14-Eotaxin** | **Combo XCIX** | **0.906** | **0.833** | **0.958** | **0.655** |
| MCP3-BLC-FGF2-TWEAK-Eotaxin | Combo C | 0.898 | 0.875 | 0.917 | 0.43 |
| **MCP3-BLC-MMP14-TWEAK-Eotaxin** | **Combo CI** | **0.908** | **0.875** | **0.958** | **0.667** |
| **MCP3-S100B-FGF2-MMP14-TWEAK** | **Combo CII** | **0.905** | **0.792** | **0.917** | **0.621** |

| | | | | | |
|---|---|---|---|---|---|
| **MCP3-S100B-FGF2-MMP14-Eotaxin** | **Combo CIII** | **0.917** | **0.875** | **0.917** | **0.586** |
| **MCP3-S100B-FGF2-TWEAK-Eotaxin** | **Combo CIV** | **0.924** | **1** | **0.708** | **0.187** |
| **MCP3-S100B-MMP14-TWEAK-Eotaxin** | **Combo CV** | **0.92** | **0.875** | **0.917** | **0.572** |
| **MCP3-FGF2-MMP14-TWEAK-Eotaxin** | **Combo CVI** | **0.91** | **0.875** | **0.958** | **0.628** |
| BLC-S100B-FGF2-MMP14-TWEAK | Combo CVII | 0.851 | 0.875 | 0.708 | 0.343 |
| BLC-S100B-FGF2-MMP14-Eotaxin | Combo CVIII | 0.884 | 0.958 | 0.75 | 0.323 |
| BLC-S100B-FGF2-TWEAK-Eotaxin | Combo CIX | 0.889 | 0.917 | 0.792 | 0.388 |
| BLC-S100B-MMP14-TWEAK-Eotaxin | Combo CX | 0.889 | 1 | 0.708 | 0.292 |
| BLC-FGF2-MMP14-TWEAK-Eotaxin | Combo CXI | 0.88 | 0.833 | 0.833 | 0.47 |
| S100B-FGF2-MMP14-TWEAK-Eotaxin | Combo CXII | 0.889 | 0.875 | 0.792 | 0.483 |
| **MCP3-BLC-S100B-FGF2-MMP14-TWEAK** | **Combo CXIII** | **0.91** | **0.875** | **0.917** | **0.471** |
| **MCP3-BLC-S100B-FGF2-MMP14-Eotaxin** | **Combo CXIV** | **0.932** | **0.875** | **0.958** | **0.598** |
| **MCP3-BLC-S100B-FGF2-TWEAK-Eotaxin** | **Combo CXV** | **0.929** | **0.875** | **0.958** | **0.565** |
| **MCP3-BLC-S100B-MMP14-TWEAK-Eotaxin** | **Combo CXVI** | **0.931** | **0.875** | **0.958** | **0.583** |
| **MCP3-BLC-FGF2-MMP14-TWEAK-Eotaxin** | **Combo CXVII** | **0.903** | **0.833** | **0.958** | **0.682** |
| **MCP3-S100B-FGF2-MMP14-TWEAK-Eotaxin** | **Combo CXVIII** | **0.913** | **0.875** | **0.917** | **0.599** |
| BLC-S100B-FGF2-MMP14-TWEAK-Eotaxin | Combo CXIX | 0.891 | 0.875 | 0.833 | 0.425 |
| **MCP3-BLC-S100B-FGF2-MMP14-TWEAK-Eotaxin** | **Combo CXX** | **0.934** | **0.875** | **0.958** | **0.62** |

Combinations with outstanding discrimination ability are in bold text. Abbreviations: AUC, area under curve; Opt, optimal; SE, sensitivity; SP, specificity

**Table S4 Combinatorial analysis of candidate biomarkers (logistic regression analysis).**

| Biomarker | AUC | SE | 95% CI | The significance level of AUC difference compared to the 7-panel AUC |
|---|---|---|---|---|
| MCP-3 | 0.862 | 0.0602 | 0.732 to 0.944 | p=0.1732 |
| BLC | 0.836 | 0.0602 | 0.701 to 0.927 | p=0.0800 |
| S100B | 0.729 | 0.074 | 0.582 to 0.847 | **p=0.0048** |
| FGF-2 | 0.731 | 0.0772 | 0.583 to 0.849 | **p=0.0071** |
| MMP-14 | 0.792 | 0.0714 | 0.650 to 0.895 | **p=0.0367** |
| TWEAK | 0.786 | 0.0665 | 0.644 to 0.891 | **p=0.0222** |
| Eotaxin | 0.752 | 0.0717 | 0.606 to 0.865 | **p=0.0094** |
| 7-biomarker panel (MCP-3/BLC/S100B/FGF-2/MMP14/TWEAK/eotaxin) | 0.953 | 0.0294 | 0.850 to 0.993 | |

Significant differences between 7-panel AUC and individual biomarker AUC are in bold text.
Abbreviations: AUC, area under curve; CI, confidence interval; SE, standard error.
